# Selective impairment of long-term depression in accumbal D1R+ MSNs involves calcium-permeable AMPARs in early Alzheimer’s disease

**DOI:** 10.64898/2026.01.19.700385

**Authors:** Nicolas Riffo-Lepe, Isaías Meza, Juliana Gonzalez-Sanmiguel, Paulina Saavedra-Sieyes, Lorena Armijo-Weingart, Armando Salinas, Loreto San Martín, Luis G. Aguayo

**Author notes:** Corresponding author Luis G. Aguayo, Ph.D. Professor, Department of Physiology University of Concepcion PO Box 160-C, (lab), Concepcion, Chile.

## Abstract

Alzheimer’s disease (AD) is increasingly associated with early circuit dysfunction preceding cognitive decline, including neuronal hyperactivity and neuropsychiatric symptoms linked to mesolimbic pathways. The nucleus accumbens (nAc), a central regulator of reward and motivational processing, exhibits early alterations in excitation/inhibition balance in patients and experimental models; however, the synaptic mechanisms underlying this vulnerability remain unclear. Here, we identify a cell-type-specific synaptic mechanism in the nAc linking early intraneuronal Aβ accumulation to circuit dysfunction. Using a double transgenic APP/PS1 mouse model expressing tdTomato in dopamine D1 receptor-positive medium spiny neurons (D1R+ MSNs), we show that long-term depression (LTD) is selectively impaired in D1R+ MSNs despite comparable Aβ levels across neuronal subtypes, revealing differential functional vulnerability. This deficit is associated with an increased contribution of calcium-permeable AMPA receptors (CP-AMPARs) and a disruption of mGluR1/5-dependent LTD, a key mechanism regulating AMPAR trafficking. Pharmacological blockade of CP-AMPARs restores synaptic depression, indicating altered receptor composition as a central feature of this phenotype. These synaptic alterations co-occur with reduced dopaminergic signaling and selective behavioral changes characterized by increased consumption of palatable reward and altered baseline context preference, while associative learning remains preserved. Together, these findings reveal a postsynaptic mechanism in which impaired mGluR-dependent plasticity permits persistent CP-AMPAR signaling, shifting synaptic balance toward increased excitatory drive and mesolimbic hyperactivity in AD.

## Introduction

Alzheimer’s disease (AD) has traditionally been conceptualized as a disorder of memory arising from cortical and hippocampal dysfunction (Selkoe and Hardy, 2016). However, neuropsychiatric symptoms, including alterations in motivation, reward processing, and impulse control, frequently emerge during early stages of disease progression and strongly predict clinical outcomes (Masters, 2015; Shah et al., 2025). Increasing clinical and experimental evidence indicates that these early manifestations are associated with network hyperactivity and disruption of excitation/inhibition (E/I) balance, with a substantial proportion of patients exhibiting epileptiform activity (Vossel et al., 2026). These observations implicate early alterations in synaptic function, yet the cellular mechanisms linking Aβ pathology to circuit hyperexcitability remain poorly defined.

A defining feature of AD pathology is the temporal dissociation between intracellular and extracellular amyloid-beta (Aβ) accumulation. Intraneuronal Aβ appears early, preceding extracellular plaque deposition, and has been shown to disrupt calcium homeostasis, synaptic function, and neuronal excitability (Alcantara-Gonzalez et al., 2025; Iulita et al., 2014; LaFerla et al., 2007). While extracellular Aβ has been extensively associated with synaptic depression and receptor loss in cortical and hippocampal circuits at advanced disease stages (Selkoe and Hardy, 2016), the synaptic consequences of endogenous intraneuronal Aβ accumulation during early, pre-plaque stages remain poorly characterized. In particular, whether intraneuronal Aβ selectively disrupts synaptic plasticity in defined neuronal populations and contributes to circuit-level hyperexcitability has not been established.

Emerging evidence indicates that early pathological changes affect the mesolimbic system, a circuit critically involved in reward processing and motivational control (Cordella et al., 2018; Fernández-Pérez et al., 2020; Kelly et al., 2021; Wu et al., 2024). The nucleus accumbens (nAc), a central integrative hub within this system, receives convergent glutamatergic and dopaminergic inputs and plays a key role in regulating hedonic drive and behavioral flexibility (Russo and Nestler, 2013). Both patients with AD and mouse models exhibit early alterations in nAc function, including changes in reward-related behaviors and neuronal excitability (Armijo-Weingart et al., 2024; Guo et al., 2022; Li et al., 2025; Nie et al., 2017), occurring prior to overt cognitive decline. Notably, intraneuronal Aβ accumulation has been reported in the nAc during pre-plaque stages (Fernández-Pérez et al., 2020), suggesting that early amyloid pathology may disrupt accumbal synaptic function. However, whether these alterations involve cell-type-specific changes in synaptic plasticity and contribute to excitation/inhibition imbalance remains unknown.

Medium spiny neurons (MSNs), the predominant neuronal population in the nAc, are inhibitory GABAergic projection neurons that provide the main output of this structure and coordinate information flow via dopamine receptor type 1 (D1R)- and dopamine receptor type 2 (D2R)-expressing pathways (Russo and Nestler, 2013; Vieitas-Gaspar et al., 2025). Balanced excitation and inhibition across these pathways is critical for appropriate motivational salience and behavioral flexibility (Li et al., 2025; Scaduto et al., 2023). Long-term depression (LTD) represents a key mechanism of synaptic plasticity in the nAc for constraining excitatory drive (Kauer and Malenka, 2007; Thomas et al., 2001). Accumbal LTD critically depends on AMPA receptor (AMPAR) remodeling driven by group I metabotropic glutamate receptor (mGluR1/5) signaling, which promotes endocytosis of calcium-permeable AMPARs (CP-AMPARs) to maintain synaptic homeostasis (Luscher and Huber, 2010; Mango and Ledonne, 2023). Disruption of this mechanism favors CP-AMPAR accumulation, enhances excitatory transmission, and shifts circuit output toward increased excitatory drive (Carr, 2020; McCutcheon et al., 2011; Wolf, 2016).

AMPARs are tetrameric ionotropic receptors composed of GluA1–4 subunits that mediate fast excitatory transmission (Henley & Wilkinson, 2016). AMPARs lacking edited GluA2, including receptors enriched in GluA1 subunits, are calcium-permeable, exhibit inward rectification, and contribute to synaptic plasticity processes involving dynamic receptor trafficking (Cull-Candy and Farrant, 2021). Although increases in CP-AMPAR expression or synaptic incorporation have been implicated in pathological conditions such as addiction, eating disorders, and epilepsy (Guo and Ma, 2021; Italia et al., 2025; Wolf, 2016), their regulation in the nAc during early AD remains unclear.

Within this framework, two key questions remain unresolved. First, are alterations in AMPAR-mediated synaptic plasticity present in the nAc during pre-plaque stages of AD in the context of intraneuronal Aβ accumulation? Second, do specific MSN subtypes exhibit differential vulnerability to these alterations? While extracellular Aβ has been shown to induce synaptic depression and AMPAR loss at advanced disease stages (Cline et al., 2018; Yang et al., 2016), the synaptic consequences of endogenous intraneuronal Aβ accumulation remain largely undefined. In particular, whether this pathology drives cell-type-specific synaptic remodeling in subcortical circuits such as the nAc is unknown.

Recent evidence indicates that dopaminergic dysfunction is an early event in AD, preceding cognitive impairment (Pilotto et al., 2025). Reduced dopamine levels, degeneration of VTA dopaminergic neurons, and decreased dopamine transporter expression in the nucleus accumbens have been reported in patients and AD mouse models (Krashia et al., 2019; Storga et al., 1996). Given the central role of dopamine in modulating excitation/inhibition balance within accumbal circuits, these findings suggest that early neuromodulatory disruption may converge with synaptic alterations to drive circuit-level dysfunction during initial stages of Aβ pathology.

However, whether intraneuronal Aβ accumulation is associated with cell-type-specific synaptic dysfunction in accumbal MSNs during pre-plaque stages of AD remains unknown. Here, we tested the hypothesis that early intraneuronal Aβ accumulation disrupts postsynaptic homeostasis and impairs long-term depression selectively in D1R+ MSNs, leading to altered excitatory balance within the nAc. Using cell-type-specific electrophysiology, synaptic plasticity assays, and measurements of calcium and dopamine signaling, we identify a selective vulnerability of accumbal D1R+ MSNs and provide mechanistic insight into the synaptic basis of early non-cognitive alterations in AD.

## MATERIALS AND METHODS

### Animals

All experimental procedures were approved by the Institutional Animal Care and Use Committee of the University of Concepción (#CEBB 1194-2022) and were conducted in accordance with national and international guidelines for the care and use of laboratory animals. Male C57BL/6J mice, double-transgenic APPswe/PS1dE9 mice (MMRRC:034832; B6.Cg-Tg(APPswe,PSEN1dE9)85Dbo/Mmjax), and Drd1a-tdTomato reporter mice (B6.Cg-Tg(Drd1a-tdTomato)6Calak/J; JAX stock #016204) were obtained from The Jackson Laboratory (Bar Harbor, ME, USA) and maintained at the Regional Center for Advanced Studies in Life Sciences (CREAV), University of Concepción. The transgenic line expresses the Swedish mutation (K594M/N595L) in amyloid precursor protein (APP) and the human presenilin-1 variant lacking exon 9 (PS1-dE9), leading to an increased Aβ production (Jankowsky et al., 2001). Drd1a-tdTomato mice express the fluorescent reporter tdTomato under the control of the dopamine D1 receptor (Drd1a) promoter, allowing selective visualization of D1R-expressing MSNs (Ade et al., 2011). To generate experimental cohorts enabling recordings from genetically fluorescent labeled D1R+ MSNs, D1RtdTomato mice were crossed with APP/PS1 mice to obtain APP/PS1;D1+tdTom offspring (Fig. 2A); WT;D1+tdTom littermates were used as controls for these experiments. Genotyping was performed according to the provider’s instructions for each line. Mice were housed in groups of 3 – 6 under a 12 h light/dark cycle with ad libitum access to food and water. Animals were used between 3 and 12 months of age. Euthanasia was performed by decapitation following anesthesia with inhaled isoflurane.

### Immunohistochemistry

Immunofluorescence experiments were done as previously reported (Armijo-Weingart et al., 2024). Briefly, mice were anesthetized with Ketamine (100 mg/kg, i.p.) and Xylazine (10 mg/kg, i.p.) and transcardially perfused with pre-warmed saline (0.9% NaCl, 35 °C), followed by freshly prepared ice-cold 4% paraformaldehyde (PFA). Brains were then dissected, post-fixed for 24 h at 4 °C, and cryoprotected in 30% sucrose for 3–5 days at 4 °C. Samples were embedded in NEG50, cooled at −20°C for 2-4 h, and stored at −80°C for at least 24 h before sectioning with a cryostat. Free-floating coronal sections (30 μm) were rinsed in Tris-phosphate buffer, permeabilized in Trisphosphate containing 1% BSA and 0.2% Triton X-100, and incubated for 24 h at 4°C with primary antibodies: MOAβ-2 (1:200, mouse, Novus Biologicals, USA) (Youmans et al., 2012), MAP2 (1:200, guinea pig, Synaptic Systems, Germany), and Iba1 (1:1000, rabbit, Alomone Labs, Germany). Sections were then incubated for 2 h with secondary antibodies (Alexa Fluor 488, Alexa Fluor 594, Alexa Fluor 647). Stained samples were mounted with DAKO fluorescent medium on glass slides and imaged using confocal microscopy at the Advanced Microscopy Center (CMA, Biobío). For each animal, at least two coronal sections were analyzed, with three distinct regions of interest (ROIs) per section. Each ROI was consistently acquired as a Z-stack (∼20 μm) for subsequent processing and quantification using FIJI and Zen software. Fluorescence intensity of MOAβ-2 was quantified by generating neuronal soma masks based on MAP2 and tdTomato signals, allowing selective measurement within identified MSN somata.

### Thioflavin-S staining

Thioflavin-S (Sigma, T1892), which binds β-sheet–rich structures present in amyloid aggregates, was used to assess extracellular amyloid plaque deposition as previously described (Fernández-Pérez et al., 2020). Coronal brain sections (30 μm) containing the nAc and hippocampus were mounted on glass slides and processed at room temperature (∼22 °C). Sections were dehydrated through a graded ethanol series (50%, 70%, 80%, 90%, 95%, and 100%; 5 min each), incubated in xylene (Winkler, XI-1670) for 10 min, and subsequently rehydrated through descending ethanol concentrations (100%, 95%, 90%, 80%, and 70%; 5 min each). Freshly prepared Thioflavin-S solution (0.05% in 50% ethanol) was filtered prior to use, and sections were incubated for 10 min protected from light. Sections were then washed in 70% ethanol (3 min) followed by distilled water (2 min), coverslipped, and stored protected from light until imaging.

Images were acquired using confocal microscopy (excitation ∼405 nm, emission collected at 430–480 nm) with a 25× oil immersion objective. Image stacks were acquired at 512 × 512 pixel resolution with a z-step of 1 μm, and acquisition parameters (laser power, gain, and offset) were kept constant across all samples. For each animal, five sections were analyzed, sampled every 100 μm, and four animals per group were included. Autofluorescence background was estimated from negative control sections processed without Thioflavin-S and subtracted from all images. Fluorescent puncta with a diameter greater than 5 μm were considered amyloid plaques based on uniform thresholding criteria. Plaque burden was quantified as [define: number or area]. Absence of detectable Thioflavin-S signal under these conditions was considered indicative of undetectable plaque deposition. All analyses were performed blinded to genotype.

### Electrophysiological recordings in coronal brain slices

Patch clamp recordings were done as previously reported (Fernández-Pérez et al., 2020). Acute coronal brain slices containing the nAc were prepared from male mice anesthetized with isoflurane and euthanized by decapitation. Brains were rapidly removed and transferred to an ice-cold, oxygenated cutting solution containing (in mM): 194 sucrose, 30 NaCl, 4.5 KCl, 1.2 NaH_2_PO_4_·H_2_O, 1 MgCl_2_·6H_2_O, 26 NaHCO_3_, and 10 Glucose (pH 7.4, equilibrated with 95% O_2_/5% CO_2_). Coronal slices (300 μm) were prepared using a vibratome (VT1200, Leica, Germany) and allowed to recover for 1 h at 32°C in artificial cerebrospinal fluid (aCSF) containing (in mM): 124 NaCl, 26 NaHCO_3_, 10 Glucose, 4.5 KCl, 2 CaCl_2_·2H_2_O, 1 MgCl_2_·6H_2_O, and 1.2 NaH_2_PO_4_·H_2_O, continuously bubbled with 95% O_2_/5% CO_2_.

Whole-cell patch-clamp recordings were performed in the nAc core using an Axopatch 200B amplifier coupled to a Digidata 1440A digitizer and pClamp 10 software (Axon Instruments). Recording pipettes (4 – 5 MΩ) were pulled from borosilicate glass capillaries (WPI) using a horizontal puller (P-1000, Sutter Instruments). During recordings, slices were continuously perfused with oxygenated aCSF at 32°C. Signals were low-pass filtered at 2 kHz and digitized at 10 kHz. Series resistance was continuously monitored and partially compensated (60–70%) throughout the recordings; cells were excluded if series resistance changed by more than 20%. To avoid pseudo-replication, only a single neuron was recorded from each brain slice. MSNs were identified based on their location within the nAc core and morphological characteristics. In experiments using Drd1a-tdTomato mice, D1R+ neurons were identified by tdTomato fluorescence, while D1R− neurons were defined by the absence of fluorescence.

### Voltage Clamp recordings

For voltage-clamp experiments, the internal pipette solution contained (in mM): 120 CsCl, 10 HEPES, 4 MgCl_2_·6H_2_O, 2 Mg-ATP, 0.5 Na_2_-GTP, and 10 BAPTA (tetra-Cs) (pH 7.4, adjusted with CsOH; 290 mOsm), together with QX-314 (1 mM) and TEA-Cl (5 mM). For rectification index experiments, Spermine (100 μM) was also included. For DHPG-induced LTD experiments, EGTA (1 mM) was used instead of BAPTA. Bath solutions were continuously perfused at a rate of 1 mL/min.

### Spontaneous synaptic currents

Spontaneous excitatory postsynaptic currents (sEPSCs) were recorded at a holding potential of −60 mV. AMPAR-mediated events were isolated by bath application of picrotoxin (PTX, 100 μM) to block GABAA and glycine receptors, and D-AP5 (50 μM) to block NMDA receptors. After break-in, cells were allowed to stabilize for at least 10 min before recording. sEPSCs were recorded under baseline conditions and subsequently in the presence of PTX and D-AP5 for a minimum of 15 min. Events were detected using a template-matching algorithm in Clampfit v11, with manual verification to exclude false positives, and at least 300 events per cell were analyzed. Parameters quantified included event frequency, amplitude, rise time, and decay time.

### Synaptic stimulation and evoked responses

Evoked excitatory postsynaptic currents (eEPSCs) were elicited using a tungsten bipolar stimulating electrode (World Precision Instruments) positioned approximately 100 μm from the recorded neuron and connected to an isolated pulse stimulator (A-M Systems). Square current pulses (1 ms, 0.05–0.5 mA) were delivered to evoke stable responses with amplitudes ≤200 pA. Stimulation intensity was adjusted to evoke stable responses and kept constant throughout each recording. Baseline recordings were initiated only after achieving stable eEPSCs that varied by no more than 30% over a period exceeding 1 min (corresponding to at least three consecutive sweeps, delivered every 20 s).

### AMPA/NMDA ratio

AMPAR- and NMDAR-mediated components were measured from eEPSCs recorded in the presence of PTX (100 μM). AMPAR responses were obtained at −60 mV. The holding potential was then shifted to +40 mV to record mixed AMPA+NMDA responses (30 sweeps, one every 20 s). D-AP5 (50 μM) was subsequently applied to isolate the AMPAR component at +40 mV. The NMDA component was calculated by subtracting the averaged AMPAR trace from the mixed response, and its amplitude was measured 20 ms after the peak of the AMPAR current. The AMPA/NMDA ratio was calculated as the peak AMPAR current divided by the NMDA current amplitude.

### Paired-pulse ratio (PPR)

Paired-pulse ratio (PPR) was assessed at −60 mV using two consecutive stimuli delivered with a 70 ms inter-stimulus interval. PPR was calculated as the ratio between the second and first eEPSC amplitudes (R2/R1), using responses ≤200 pA.

### Rectification index (RI)

Rectification properties of AMPAR-mediated currents were assessed using CsCl-based internal solution containing spermine (100 μM), in the presence of PTX (100 μM) and D-AP5 (50 μM). eEPSCs were recorded at holding potentials ranging from −60 to +40 mV in 20 mV increments. For each potential, 30 sweeps were collected and averaged. The rectification index was calculated as the ratio of the absolute current amplitude at +40 mV to that at −60 mV.

### NASPM sensitivity

To assess the contribution of calcium-permeable AMPARs, MSNs were voltage-clamped at −60 mV. After establishing a stable baseline for at least 10 min (one stimulus every 20 s), NASPM (150 μM) was bath-applied for a minimum of 15 min. Inhibition was expressed as the percentage reduction in mean eEPSC amplitude, comparing baseline responses with those recorded during the last 5 min of NASPM application.

### Long-term depression

All LTD experiments were performed in the presence of PTX (100 μM) and D-AP5 (50 μM). For HFS-induced LTD, recordings were obtained with CsCl-based internal solution. After recording a stable baseline for at least 10 min, LTD was induced using four trains of stimuli delivered at 100 Hz (1 ms pulses), separated by 20 s. eEPSCs were recorded for at least 40 min following induction. Responses were normalized to baseline before quantification and LTD magnitude was calculated as the percentage change in normalized eEPSC amplitude during the last 5 min relative to baseline.

For mGluR1/5-dependent LTD, recordings were performed using internal solution containing EGTA (1 mM). After a stable 10 min baseline, (RS)-3,5-dihydroxyphenylglycine (DHPG, 50 μM) was bath-applied for 5 min, and eEPSCs were monitored for at least 25 min thereafter. LTD magnitude was calculated as the percentage reduction in mean eEPSC amplitude during the final 5 min relative to baseline.

### Stereotaxic injections

Five-month-old male WT and APP/PS1 mice were used. Stereotaxic surgery was performed to deliver adeno-associated viruses (AAVs) expressing the genetically encoded calcium indicator GCaMP6s under the synapsin promoter, as previously described (Armijo-Weingart et al., 2024). A total of 200 nL of AAV1-Syn-GCaMP6s.WPRE.SV40 (1.76 × 10^13^ GC/ml; Addgene #100843-AAV1) or 400 nL of pAAV-CAG-dLight1.1 (7 × 10¹² vg/ml; Addgene #111067-AAV5) was injected bilaterally into the nAc using a stereotaxic alignment system (Kopf Instruments) at 50 nL per minute. Injection coordinates relative to bregma were: AP +0.13 mm, ML ±0.11 mm, and DV −0.4 mm (Allen Brain Atlas). Animals received post-operative analgesia (Meloxicam 2 mg/kg). Mice were anesthetized with 4% isoflurane/oxygen and positioned in a stereotaxic frame; anesthesia was maintained with 2–3% isoflurane/oxygen throughout the procedure. After leveling the skull, a small craniotomy was made at the target site. A 1 μL Neuros Hamilton syringe was lowered slowly to the desired depth, and viral solution was delivered. The syringe was left in place for 3 min post-infusion before withdrawal, and incisions were closed with Leukosan adhesive.

### Calcium and Dopamine photometry

Brain slice photometry were evaluated as previously reported (Salinas et al., 2023). Two to three weeks after AAV injection, mice were 6 months old at the time of experiments. Acute coronal slices (300 μm) containing the nAc were prepared for calcium or dopamine imaging. Slices were transferred to an upright microscope and continuously perfused with oxygenated aCSF (1 mL/min). The recording region of interest (medial to the anterior commissure, corresponding to the nAc core) was visualized under fluorescence to confirm GCaMP6s or dLight1.1 expression. A bipolar stimulating electrode (DS3 Isolated Current Stimulator, Digitimer, UK) was placed on the slice surface near the area of interest. Stimulation consisted of single electrical pulses (400–800 μA, 1 ms duration) delivered every 2 min, generating one evoked fluorescence transient per stimulus. Transients were measured by slice photometry using a Horiba PTI D-104 Microscope Photometer equipped with a 710 nm photomultiplier tube mounted on an Olympus BX51 microscope, with a 120 LED Boost High-Power illumination system and appropriate excitation/emission filters (488 nm for GCaMP6s and 405 nm for dLight1.1).

Fluorescence signals were acquired using PatchMaster software and expressed as ΔF/F_0_, where F_0_ was defined as the mean baseline fluorescence prior to stimulation. A stable baseline was recorded for 12 min (six responses), followed by bath application of the AMPAR antagonist CNQX and continued stimulation for an additional 12 min until responses reached a plateau. For analysis, ΔF/F_0_ values were normalized to baseline and expressed as percentage change. Drug effects were quantified by comparing the mean response during the last 6 min in the presence of CNQX with baseline for each slice. At least two recordings per mouse were obtained from independent nAc slices; each slice was treated as an individual observation, while the number of animals contributing to each dataset is reported in the corresponding figure legends.

### Western blot

Protein expression analysis were done as previously described (Armijo-Weingart et al., 2024). The nucleus accumbens (nAc) was microdissected from coronal brain slices obtained from 6-month-old WT and APP/PS1 male mice. Each biological replicate corresponded to the entire nAc isolated from a single animal. Tissue samples were homogenized in ice-cold RIPA buffer supplemented with protease and phosphatase inhibitor cocktails. Lysates were centrifuged at 14,000 × g for 15 min at 4 °C, and supernatants were collected for protein quantification. Equal amounts of protein (50 μg per lane) were denatured in buffer, separated by SDS–PAGE at constant 80 Volt for 3 hours, and transferred onto PVDF membranes at 250 mA for 3 hours. Membranes were blocked in 5% non-fat dry milk prepared in TBS-T (Tris-buffered saline containing 0.1% Tween-20) for 1 h at room temperature and then incubated overnight at 4 °C with primary antibodies against SV2 (DSHB #AB2315387), PSD95 (Synaptic Systems #124011), GluA1 (Synaptic Systems #182011), GluA2 (Synaptic Systems #182103), and Gβ (Santa Cruz, sc-166123) (used as loading control).

### qRT-PCR

The nAc was microdissected from 300 μm coronal brain slices. Total RNA was extracted using TRIzol reagent according to the manufacturer’s instructions and treated with DNase to eliminate potential genomic DNA contamination. Complementary DNA (cDNA) was synthesized from 2 μg of total RNA using reverse transcriptase and oligo(dT) primers. Quantitative real-time PCR was performed for 40 cycles using SYBR Green Universal Master Mix (Agilent Technologies) and gene-specific primers targeting NMDA receptor subunits and AMPA receptor subunits. The following primer pairs were used: Gria1 (GluA1), forward 5′-ACCCTCCATGTGATCGAAATG-3′ and reverse 5′-GGTTCTATTCTGGACGCTTGAG-3′; Gria2 (GluA2), forward 5′-AAAGAATACCCTGGAGCACAC-3′ and reverse 5′-CCAAACAATCTCCTGCATTTCC-3′; Grin1 (NMDA receptor subunit 1), forward 5′-AAATGTGTCCCTGTCCATACTC-3′ and reverse 5′-CCTGCCATGTTCTCAAAAGTG-3′; Grin2b (NMDA receptor subunit 2B), forward 5′-GAACGAGACTGACCCAAAGAG-3′ and reverse 5′-CAGAAGCTTGCTGTTCAATGG-3′. Cyclophilin A was used as the housekeeping gene, with forward primer 5′-ATAATGGCACTGGTGGCAAGTC-3′ and reverse primer 5′-ATTCCTGGACCCAAAACGCTCC-3′. Relative expression levels were normalized to Cyclophilin A and calculated using the 2^-ΔΔCt^ method.

### Statistical analysis

Animals were randomly selected for recordings without prior knowledge of their APP/PS1 or WT genotype; only tdTomato fluorescence was used to identify D1R-expressing MSNs when applicable. Data acquisition and analysis were performed blind to genotype. The number of cells and animals analyzed for each experiment is reported in the corresponding figure legends. Individual neurons were treated as independent observations unless otherwise specified, and data were derived from multiple animals per group.

All electrophysiological data were analyzed using Clampfit v11 (Molecular Devices). Synaptic event detection was performed using a template-matching algorithm implemented in Clampfit, with manual verification to exclude false positives. Data were organized in Microsoft Excel and subsequently imported into GraphPad Prism (v10) for statistical analyses and figure preparation. Normality was assessed using the Shapiro–Wilk test. For comparisons between two independent groups, unpaired two-tailed Student’s t-tests or Welch’s t-tests were used when variance was unequal, whereas Mann–Whitney U tests were applied for non-normally distributed data. For experiments involving repeated measures over time, LTD time-course analyses, mixed-effects models with restricted maximum likelihood (REML) estimation were used, with Genotype, Cell type, or Age as fixed factors and animal identity included as a random effect. When appropriate, post hoc comparisons were performed using Sidak’s or Tukey’s multiple-comparison tests. Paired-pulse ratio analyses were assessed using unpaired two-tailed t-tests. Data are presented as mean ± s.e.m., and statistical significance was defined as p < 0.05. Exact p values, test statistics, degrees of freedom, and sample sizes are reported in the corresponding figure legends.

## RESULTS

### Intraneuronal Aβ accumulation in the nucleus accumbens defines a pre-plaque stage and associates with selective alterations in excitatory synaptic transmission

To define the temporal stage of amyloid pathology in the nAc, we first assessed Aβ accumulation and plaque deposition across ages in the murine AD models. Intraneuronal Aβ was detected in accumbal neurons of APP/PS1 mice as early as 3 months of age, with increased signal intensity compared with WT controls (Fig. S1A, B). At later stages, extracellular plaques remained largely absent in the nAc at 6 and 9 months and were only sparsely detected at 12 months, whereas cortical regions exhibited earlier and more prominent plaque deposition (Fig. S1C). In addition, no differences were observed in the number or fluorescence intensity of Iba1-positive microglia at 6 months (Fig. S1D F), indicating the absence of overt neuroinflammatory changes. Together, these findings define a pre-plaque stage in the nAc characterized by predominantly intraneuronal Aβ accumulation.

We next examined whether intraneuronal Aβ accumulation is differentially distributed across MSN subtypes and whether it is associated with alterations in synaptic function. In 6-month-old APP/PS1;D1-tdTom mice, Aβ immunoreactivity was prominently detected within neuronal somata, with no evidence of extracellular plaque deposition (Fig. 1A,B). Quantitative analysis revealed increased Aβ levels in both D1R+ and D1R− MSNs compared with WT controls (Fig. 1C), indicating that intraneuronal Aβ accumulation is broadly distributed across MSN subtypes.

**Figure 1.**
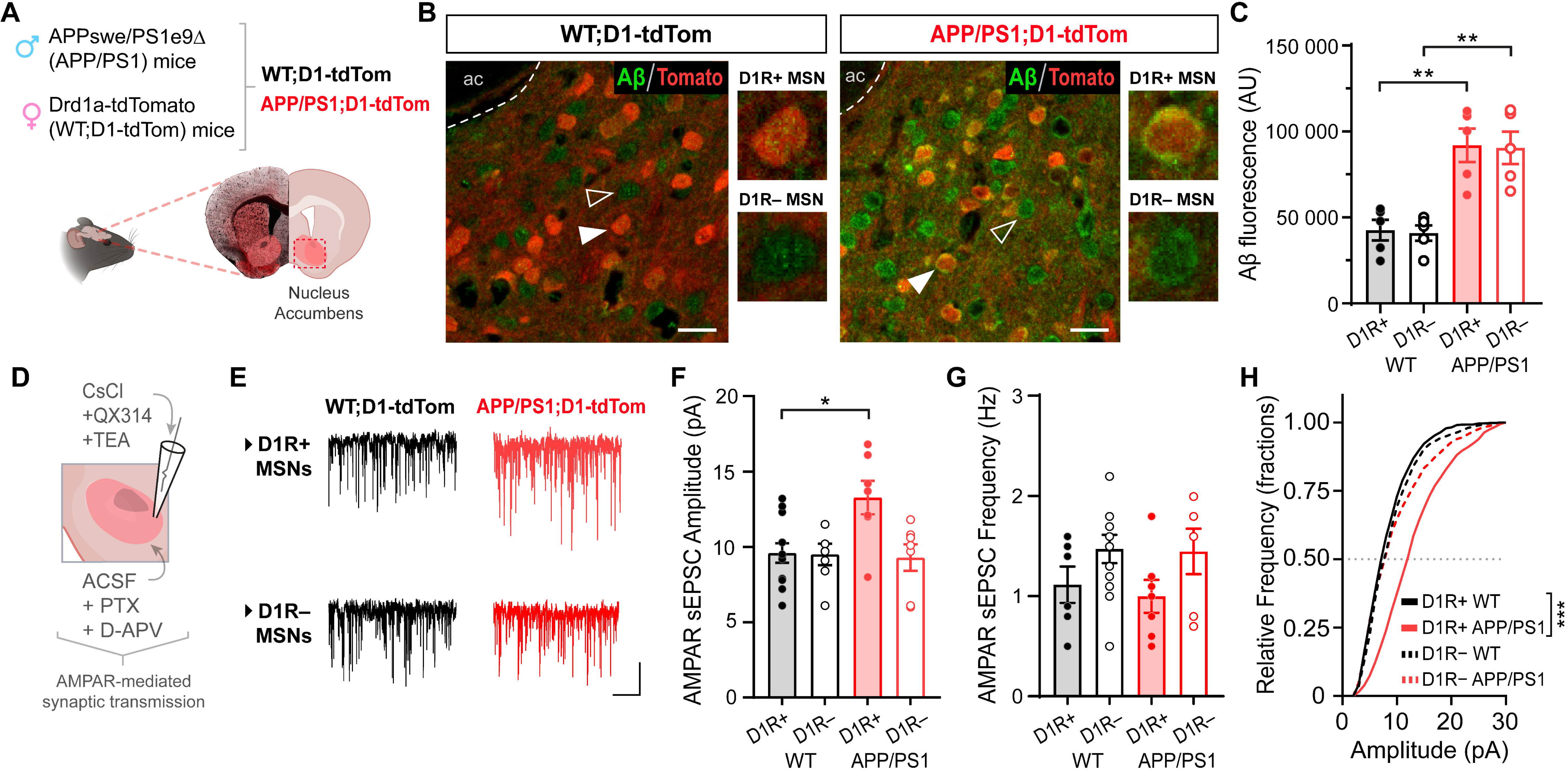
Intraneuronal Aβ accumulation and altered excitatory synaptic transmission in accumbal MSNs during pre-plaque stages. (A) Schematic representation of the experimental strategy. Crossing APP/PS1 mice with Drd1a-tdTomato reporter mice enables identification of dopamine D1 receptor-expressing (D1R+) MSNs. (B) Representative confocal images of nucleus accumbens (nAc) sections from 6-month-old WT;D1-tdTom and APP/PS1;D1-tdTom mice. Sections were labeled for Aβ (MOAβ-2, green) and tdTomato (red). Filled arrowheads indicate D1R+ MSNs (tdTomato+), whereas open arrowheads denote D1R− MSNs (tdTomato−). Right small panels show single-cell magnifications corresponding to each MSN subtype. Scale bars: 20 μm. (C) Quantification of intraneuronal Aβ fluorescence intensity in D1R+ and D1R− MSN somas at 6 months. APP/PS1 mice exhibit increased Aβ levels in both MSN subtypes compared with WT (one-way ANOVA followed by Tukey’s multiple comparisons test). Each point represents the mean value per animal (n = 5 per genotype). Total cells analyzed: WT, 50 D1R+ and 47 D1R−; APP/PS1, 40 D1R+ and 45 D1R− MSNs. (D) Schematic representation of whole-cell voltage-clamp recordings used to assess AMPAR-mediated spontaneous excitatory postsynaptic currents (sEPSCs) in accumbal MSNs. (E) Representative sEPSC traces recorded at −60 mV from D1R+ (top) and D1R− (bottom) MSNs in WT;D1-tdTom (black) and APP/PS1;D1-tdTom (red) mice. Scale bars: 6 pA, 5 s. (F) Summary of sEPSC amplitude showing a significant interaction between genotype and MSN subtype (two-way ANOVA followed by Sidak multiple comparisons test). Post hoc comparisons revealed increased sEPSC amplitude in APP/PS1 D1R+ MSNs compared with WT D1R+ MSNs, with no differences in D1R− MSNs. (G) Quantification of sEPSC frequency shows no significant differences across groups (one-way ANOVA, p = 0.155). (H) Cumulative probability distributions of sEPSC amplitudes showing a rightward shift selectively in APP/PS1 D1R+ MSNs compared with WT D1R+ MSNs, with no differences observed between WT and APP/PS1 D1R− MSNs (Kolmogorov–Smirnov test).

Given this widespread Aβ distribution, we next assessed AMPAR-mediated synaptic transmission in identified MSN populations. Whole-cell recordings revealed a selective increase in sEPSC amplitude in D1R+ MSNs from APP/PS1 mice, whereas no differences were observed in D1R− MSNs (Fig. 1D–F). The sEPSC frequency remained unchanged across genotypes and MSN subtypes (Fig. 1G). Analysis of amplitude distributions further showed a rightward shift selectively in D1R+ MSNs (Fig. 1H), consistent with a postsynaptic increase in excitatory synaptic strength.

Importantly, these alterations were not present at earlier stages. Analysis of AMPAR-mediated sEPSCs in 3-month-old mice revealed no differences between WT and APP/PS1 MSNs in amplitude, frequency, or kinetic properties (Fig. S2A–I). In contrast, at 6 months, APP/PS1 MSNs exhibited increased sEPSC amplitude without changes in frequency or kinetics (Fig. S2J–R), indicating that functional alterations in excitatory synaptic transmission emerge during the pre-plaque stage.

Together, these results indicate that early intraneuronal Aβ accumulation in the nAc occurs in the absence of extracellular pathology and neuroinflammation, associated with selective alterations in AMPAR-mediated synaptic transmission in D1R+ MSNs despite comparable Aβ levels across MSN subtypes.

### Selective impairment of long-term depression in the nucleus accumbens emerges at pre-plaque stages and selectively affects D1R+ MSNs

To determine whether early intraneuronal Aβ accumulation is associated with alterations in synaptic plasticity, we evaluated long-term depression (LTD) in accumbal MSNs from WT and APP/PS1 mice across ages. At 3 months, HFS-induced LTD was robust and comparable between genotypes. In contrast, at 6 months, LTD magnitude was reduced in APP/PS1 mice compared with WT controls (Fig. S3A–G), indicating that impairments in synaptic plasticity emerge during pre-plaque stages. Consistent with this, mGluR1/5-dependent LTD induced by DHPG was also attenuated in APP/PS1 MSNs (Fig. S3H–K). Paired-pulse ratio (PPR) remained unchanged between genotypes (Fig. S3L–N), suggesting preserved presynaptic release probability and supporting a postsynaptic origin of LTD impairment.

We next examined whether LTD disruption is cell-type specific. Whole-cell recordings in D1R+ and D1R− MSNs from WT;D1-tdTom and APP/PS1;D1-tdTom mice revealed a selective reduction of HFS-induced LTD in D1R+ MSNs from APP/PS1 mice, whereas D1R− MSNs exhibited preserved LTD across genotypes (Fig. 2A–D). Consistent with previous observations, PPR remained unchanged across MSN subtypes (Fig. 2E). Together, these results indicate that synaptic plasticity deficits emerge at pre-plaque stages and selectively affect D1R+ MSNs, consistent with a postsynaptic mechanism underlying LTD impairment.

**Figure 2.**
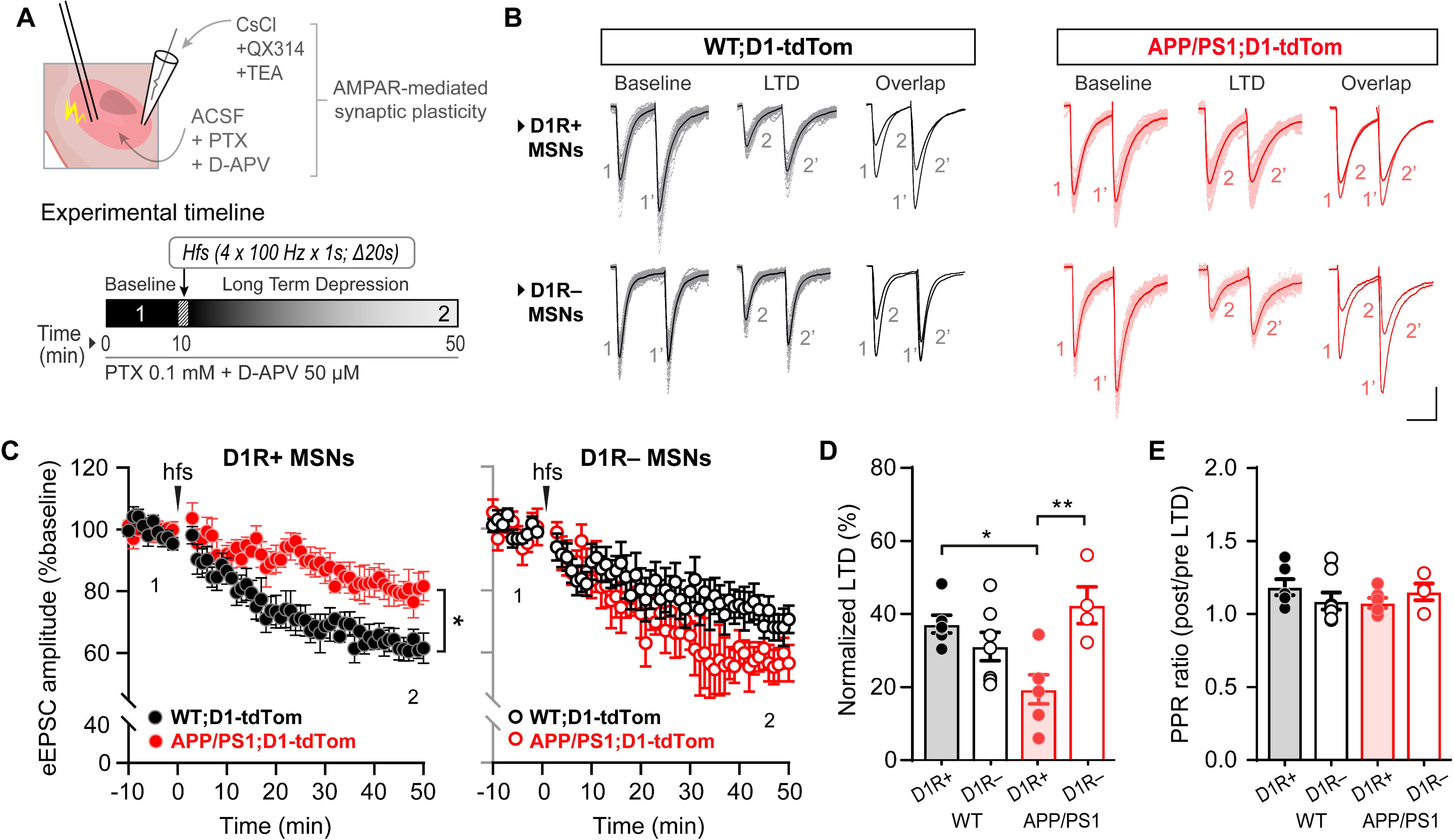
HFS-induced LTD is selectively impaired in D1R+ MSNs, but preserved in D1R− MSNs, in the nAc of 6-month-old APP/PS1 mice. (A) Schematic representation of the experimental configuration for whole-cell voltage-clamp recordings of AMPAR-mediated synaptic transmission in accumbal MSNs. The experimental timeline indicates baseline eEPSC acquisition (1; 10 min), induction of LTD using high-frequency stimulation (HFS; four trains at 100 Hz, 1 s duration, delivered every 20 s), and subsequent monitoring of LTD expression. LTD magnitude was calculated by normalizing the mean eEPSC amplitude during the final 10 min of recording (2; 40–50 min) to baseline. (B) Representative paired-pulse evoked EPSC traces recorded from D1R+ and D1R− MSNs in WT;D1-tdTom (black) and APP/PS1;D1-tdTom (red) mice before (1, baseline) and after HFS-induced LTD (2). Overlays illustrate the reduction in eEPSC amplitude following LTD induction. Scale bars: 50 pA, 35 ms. (C) Time course of normalized eEPSC amplitude in D1R+ (left) and D1R− (right) MSNs from WT and APP/PS1 mice. (D) Quantification of LTD magnitude calculated from the final 10 min of recording (40–50 min). LTD was reduced in APP/PS1 D1R+ MSNs compared with WT D1R+ MSNs, whereas no differences were observed in D1R− MSNs (one-way ANOVA followed by Tukey multiple comparisons test). LTD magnitude also differed between D1R+ and D1R− MSNs within APP/PS1 mice. (E) Quantification of paired-pulse ratio (PPR; post/pre) showing no significant differences across genotypes or MSN subtypes (Kruskal–Wallis test followed by Dunn multiple comparisons test). Each data point represents a single recorded neuron. Number of neurons/mice: WT D1R+ (n = 6/5), WT D1R− (n = 7/5), APP/PS1 D1R+ (n = 6/5), APP/PS1 D1R− (n = 4/4). Data are presented as mean ± SEM. *p < 0.05, **p < 0.01.

### Alterations in AMPAR-mediated synaptic transmission in the nucleus accumbens are not explained by transcriptional changes

To further characterize the synaptic alterations underlying LTD impairment, we evaluated AMPAR- and NMDAR-mediated synaptic transmission in accumbal MSNs from 6-month-old WT and APP/PS1 mice. Whole-cell recordings at +40 mV revealed a reduction in the AMPA/NMDA ratio in APP/PS1 mice compared with WT controls (Fig. 3A–C), indicating a relative shift in excitatory synaptic composition.

**Figure 3.**
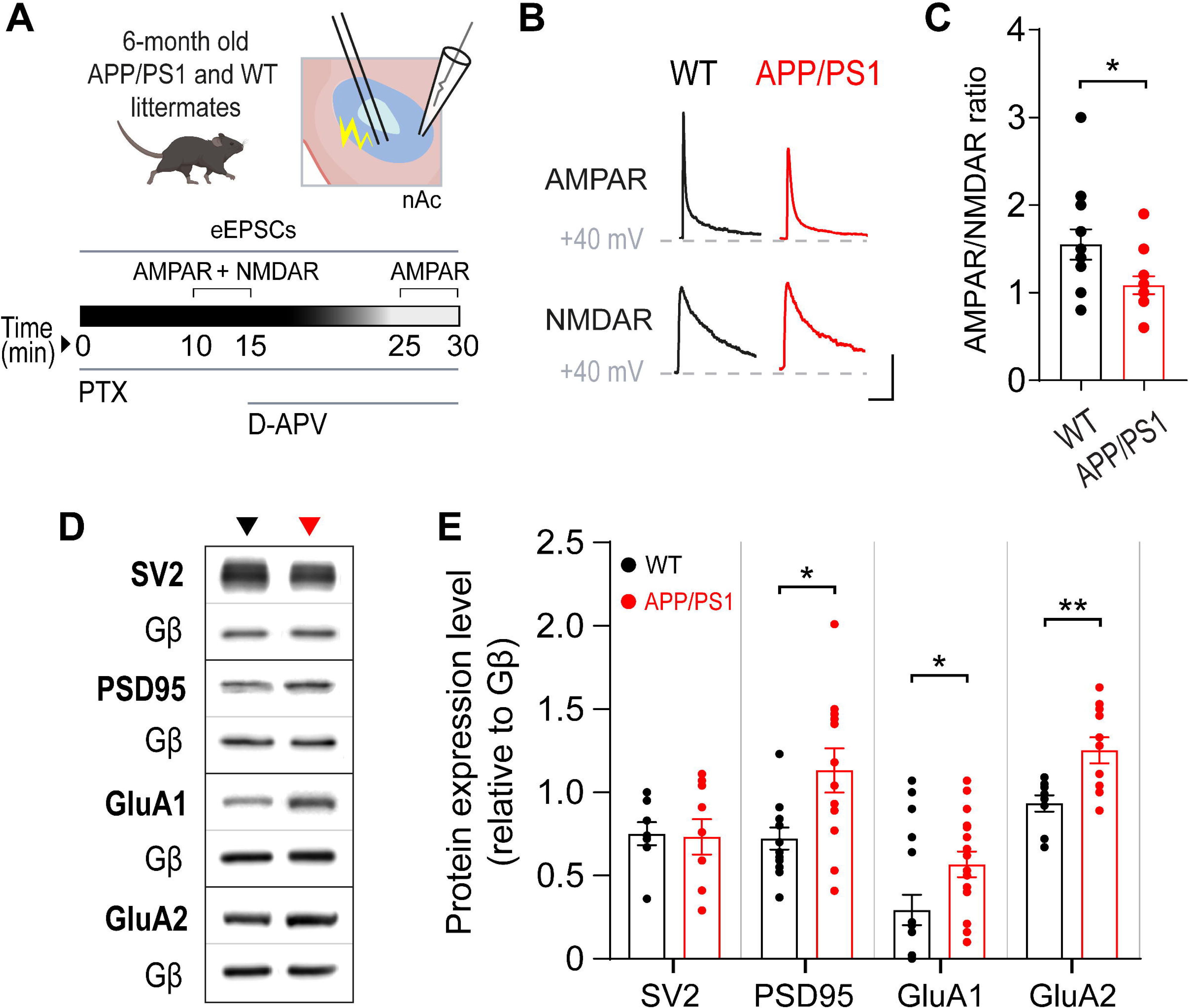
Reduction of AMPA/NMDA ratio and altered AMPAR expression in the nucleus accumbens of APP/PS1 mice. (A) Schematic representation of the experimental design used to assess AMPAR- and NMDAR-mediated synaptic transmission in accumbal MSNs. Whole-cell voltage-clamp recordings were performed at +40 mV to obtain mixed AMPA+NMDA eEPSCs, followed by pharmacological isolation of AMPAR-mediated currents after D-APV application. The NMDAR-mediated component was obtained by digital subtraction. (B) Representative AMPAR- and NMDAR-mediated eEPSCs recorded at +40 mV from MSNs of WT (black) and APP/PS1 (red) mice. Scale bars: 50 pA, 100 ms. (C) Quantification of the AMPA/NMDA ratio showing a reduction in APP/PS1 mice compared with WT (unpaired two-tailed t-test). n = 12/4 (WT) and 13/4 (APP/PS1) neurons/mice. (D) Representative Western blots of synaptic proteins extracted from the nucleus accumbens of WT and APP/PS1 mice. (E) Quantification of protein expression levels normalized to Gβ. No differences were observed in the presynaptic marker SV2, whereas PSD95, GluA1, and GluA2 levels were increased in APP/PS1 mice (unpaired two-tailed t-test, Welch’s correction, or Mann–Whitney test as appropriate). n values are indicated in the figure. Data are presented as mean ± SEM. *p < 0.05, **p < 0.01.

To determine whether these functional changes are associated with alterations in synaptic protein expression, we quantified key glutamatergic markers in the nAc. While levels of the presynaptic protein SV2 were unchanged, postsynaptic markers showed selective alterations. PSD95 expression was increased in APP/PS1 mice, consistent with changes in postsynaptic scaffolding. In addition, GluA1 and GluA2 subunits were elevated in APP/PS1 mice compared with WT controls (Fig. 3D–E), indicating altered AMPAR composition at the protein level.

To assess whether these changes are driven by transcriptional regulation, we measured mRNA expression of glutamate receptor subunits in the nAc. No differences were observed between WT and APP/PS1 mice for Gria1, Gria2, Grin1, or Grin2b at any age analyzed (Fig. S4A–D), indicating that the observed synaptic alterations are not explained by changes in gene expression.

Changes in the AMPA/NMDA ratio may reflect alterations in receptor composition or conductance properties, rather than a simple reduction in AMPAR-mediated transmission. In particular, an increased contribution of calcium-permeable AMPARs, which exhibit inward rectification and reduced current at depolarized potentials, could contribute to the observed decrease in AMPA/NMDA ratio (Cull-Candy and Farrant, 2021). Consistent with this, the upregulation of AMPAR subunits observed in APP/PS1 mice aligns with the LTD deficit described above, as increased AMPAR abundance is typically associated with enhanced synaptic strength rather than synaptic depression (Italia et al., 2025; Muñoz de Leon-Lopez et al., 2025). Together, these findings are compatible with altered AMPAR composition and suggest aberrant receptor remodeling in the nAc during early stages of APP/PS1 pathology.

### Increased contribution of calcium-permeable AMPARs to excitatory synaptic transmission in the nucleus accumbens of APP/PS1 mice

To determine whether the alterations in AMPAR-mediated transmission observed in APP/PS1 mice are associated with changes in AMPAR functional properties, we assessed AMPAR-dependent Ca^2+^ signaling and rectification properties in accumbal MSNs. GCaMP6s imaging revealed that blockade of AMPARs with CNQX produced a greater reduction in evoked Ca^2+^ transients in APP/PS1 slices compared with WT controls (Fig. 4A–D), indicating an increased contribution of AMPAR-mediated Ca^2+^ influx. Consistent with this, recordings across holding potentials showed reduced current amplitudes at depolarized potentials in APP/PS1 MSNs, resulting in a lower rectification index compared with WT (Fig. 4E–H). This inwardly rectifying behavior is characteristic of GluA2-lacking AMPARs, which are subject to voltage-dependent polyamine block (Cull-Candy and Farrant, 2021)

**Figure 4.**
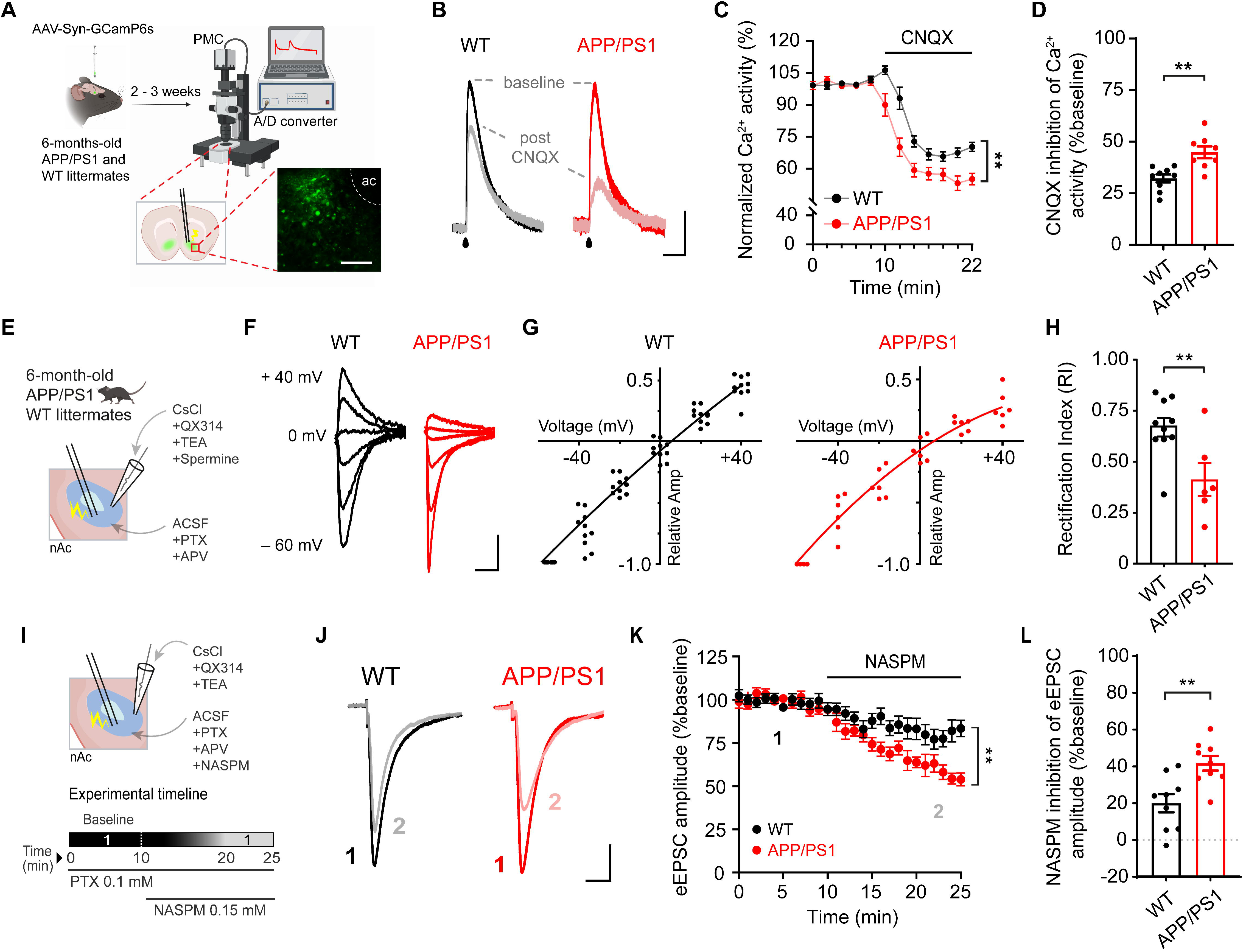
Increased contribution of calcium-permeable AMPARs in the nucleus accumbens of APP/PS1 mice at pre-plaque stages. (A) AAV-Syn-GCaMP6s was injected into the nucleus accumbens (nAc) of WT and APP/PS1 mice and allowed to express for 2–3 weeks prior to acute slice preparation for calcium imaging. (B) Representative electrically evoked calcium transients recorded in nAc slices from 6-month-old WT (black) and APP/PS1 (red) mice before and after application of the AMPAR antagonist CNQX. Scale bars: 25% ΔF/F, 2 s. (C) Time course of normalized GCaMP6s fluorescence (percentage of baseline) showing a greater reduction following CNQX application in APP/PS1 slices. (D) Quantification of CNQX-sensitive calcium activity showing increased inhibition in APP/PS1 mice compared with WT (unpaired two-tailed t-test). n = 9/4 (WT) and 8/4 (APP/PS1) recordings/mice. (E) Schematic representation of whole-cell voltage-clamp recordings used to assess AMPAR-mediated eEPSCs in accumbal MSNs. Recordings were performed with intracellular spermine (100 μM). (F) Representative eEPSC traces recorded at holding potentials from −60 to +40 mV (20 mV increments) from WT (black) and APP/PS1 (red) MSNs. Scale bars: 50 pA, 15 ms. (G) Current–voltage relationships showing reduced inward rectification in APP/PS1 MSNs compared with WT. (H) Quantification of the rectification index (RI) showing a decrease in APP/PS1 MSNs (unpaired two-tailed t-test). n = 10/4 (WT) and 6/4 (APP/PS1) neurons/mice. (I) Experimental protocol for pharmacological isolation of calcium-permeable AMPARs using NASPM. (J) Representative eEPSC traces recorded before and after NASPM application (150 μM). Scale bars: 30 pA, 25 ms. (K) Time course of normalized eEPSC amplitude (percentage of baseline) showing greater NASPM-induced inhibition in APP/PS1 MSNs. (L) Quantification of NASPM-sensitive eEPSC inhibition showing increased sensitivity in APP/PS1 MSNs compared with WT (unpaired two-tailed t-test). n = 9/4 (WT) and 9/4 (APP/PS1) neurons/mice. Data are presented as mean ± SEM. Each data point represents a single recording or neuron. *p < 0.05, **p < 0.01.

To directly assess the functional contribution of calcium-permeable AMPARs, we applied the selective antagonist NASPM, which induced a greater reduction in eEPSC amplitude in APP/PS1 MSNs compared with WT neurons (Fig. 4I–L), indicating an increased contribution of NASPM-sensitive AMPAR components to synaptic transmission.

Together, these findings indicate that early alterations in AMPAR-mediated synaptic transmission in the nAc of APP/PS1 mice are accompanied by changes in receptor functional properties consistent with an increased contribution of calcium-permeable AMPARs. These alterations provide a basis to examine whether aberrant AMPAR composition contributes to the impairment of synaptic plasticity mechanisms observed at this stage.

### mGluR-dependent LTD is selectively impaired in D1R+ MSNs and is associated with persistent CP-AMPAR contribution in APP/PS1 mice

Given that APP/PS1 mice exhibit both impaired LTD and increased contribution of CP-AMPARs in the nAc, we next examined whether these alterations are associated with changes in mGluR1/5-dependent LTD, a form of synaptic plasticity that promotes AMPAR endocytosis (Mango and Ledonne, 2023). Whole-cell recordings were performed in D1R+ and D1R− MSNs from 6-month-old WT;D1-tdTom and APP/PS1;D1-tdTom mice using a sequential protocol consisting of baseline acquisition, DHPG-induced LTD, and subsequent application of the CP-AMPAR antagonist NASPM (Fig. 5A,B).

**Figure 5:**
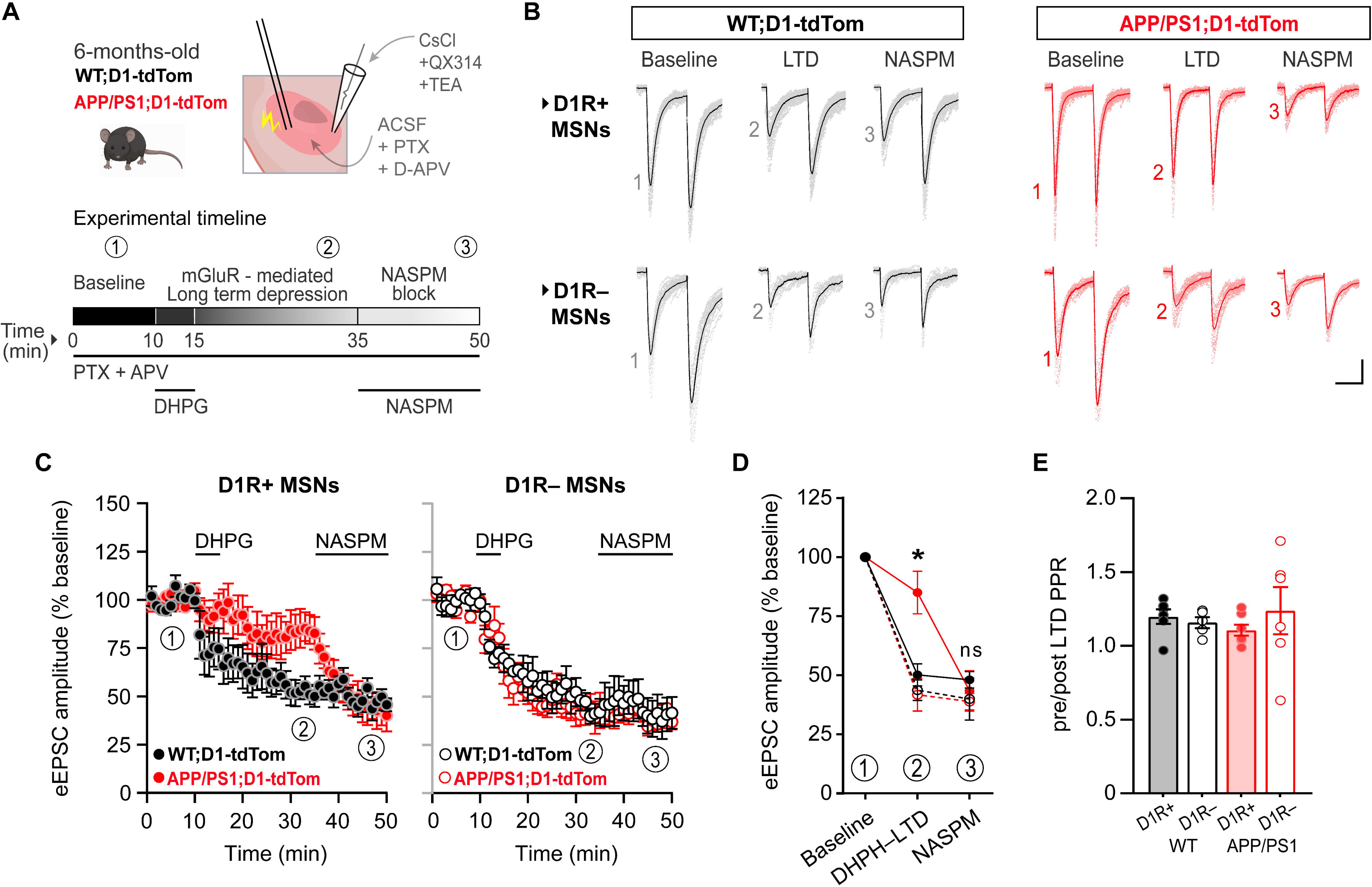
mGluR-dependent LTD is selectively impaired in D1R+ MSNs and partially restored by NASPM in APP/PS1 mice. (A) Schematic representation of whole-cell voltage-clamp recordings in accumbal MSNs from 6-month-old WT;D1-tdTom and APP/PS1;D1-tdTom mice. After baseline acquisition (1), LTD was induced by bath application of DHPG (50 μM, 5 min) and quantified 30–35 min post-application (2). NASPM (150 μM) was subsequently applied, and its effect was measured during the final 5 min of recording (3). (B) Representative AMPAR-mediated eEPSC traces recorded from D1R+ (top) and D1R− (bottom) MSNs at baseline, after DHPG-induced LTD, and during NASPM application in WT (black) and APP/PS1 (red) mice. Scale bars: 50 pA, 40 ms. (C) Time course of normalized eEPSC amplitude (percentage of baseline) in D1R+ (left) and D1R− (right) MSNs. DHPG-induced LTD was reduced in D1R+ MSNs from APP/PS1 mice and partially recovered following NASPM application, whereas D1R− MSNs showed comparable LTD between genotypes. Horizontal bars indicate periods of DHPG and NASPM application. (D) Quantification of normalized eEPSC amplitudes at baseline, after DHPG-induced LTD, and during NASPM application. LTD was reduced in APP/PS1 D1R+ MSNs compared with WT D1R+ MSNs and APP/PS1 D1R− MSNs (mixed-effects model with Tukey multiple comparisons test). (E) Quantification of paired-pulse ratio (PPR; post/pre LTD) showing no significant differences across genotypes or MSN subtypes (Kruskal–Wallis test followed by Dunn multiple comparisons test). Each data point represents a single recorded neuron. Number of neurons/mice: WT D1R+ (n = 6/4), WT D1R− (n = 5/4), APP/PS1 D1R+ (n = 7/5), APP/PS1 D1R− (n = 6/5). Data are presented as mean ± SEM.

In WT D1R+ MSNs, DHPG induced a robust and sustained depression of eEPSC amplitude (Fig. 5C,D). Under these conditions, subsequent NASPM application did not further reduce synaptic responses, indicating a minimal contribution of CP-AMPARs following LTD expression. In contrast, in APP/PS1 D1R+ MSNs, DHPG failed to induce significant synaptic depression (Fig. 5C,D). Notably, subsequent NASPM application produced a marked reduction in eEPSC amplitude, restoring synaptic depression to levels comparable to those observed in WT neurons after LTD induction (Fig. 5C,D). Paired-pulse ratio remained unchanged across conditions (Fig. 5E), indicating preserved presynaptic release probability.

Together, these findings indicate that mGluR1/5-dependent LTD is selectively impaired in D1R+ MSNs of APP/PS1 mice and is associated with a persistent functional contribution of CP-AMPARs. Acute blockade of CP-AMPARs restores synaptic depression under these conditions, consistent with altered AMPAR remodeling during early stages of pathology.

### Selective alterations in dopaminergic signaling and reward-related behavior in APP/PS1 mice at pre-plaque stages

To determine whether early synaptic alterations in the nAc of APP/PS1 mice are accompanied by changes in dopaminergic signaling, dopamine dynamics were assessed using the genetically encoded fluorescent sensor dLight1.1 expressed in the nAc (Fig. 6A). Acute slice photometry revealed that electrical stimulation evoked robust dopamine-dependent fluorescence transients in WT mice, whereas responses were reduced in APP/PS1 mice (Fig. 6B). Consistently, input–output analysis showed a decrease in normalized dLight1.1 activity across stimulation intensities in APP/PS1 mice compared with WT controls (Fig. 6C), indicating reduced evoked dopaminergic signaling in the nAc at pre-plaque stages.

**Figure 6:**
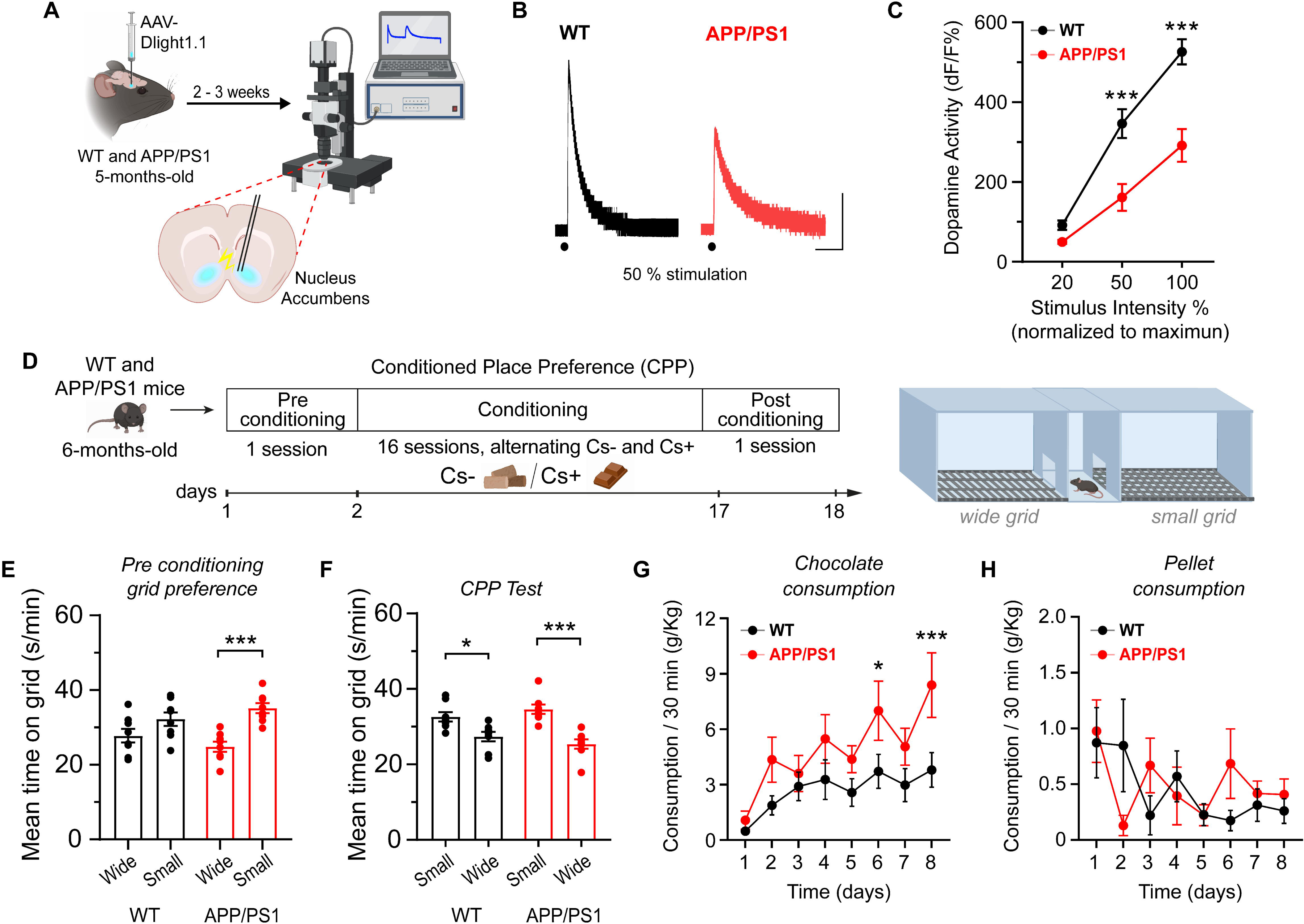
Reduced dopamine signaling in the nucleus accumbens and altered reward-related behavior in APP/PS1 mice. (A) AAV-dLight1.1 was injected into the nucleus accumbens (nAc) of WT and APP/PS1 mice and allowed to express for 2–3 weeks prior to acute slice preparation for dopamine imaging. Dopamine-dependent fluorescence signals were recorded using slice photometry following electrical stimulation. (B) Representative dLight1.1 fluorescence traces evoked by electrical stimulation in nAc slices from WT (black) and APP/PS1 (red) mice. Scale bars: 100% ΔF/F, 5 s. (C) Input–output relationship between dLight1.1 fluorescence (ΔF/F, normalized to maximum) and stimulus intensity showing reduced dopamine signals in APP/PS1 mice (two-way ANOVA followed by Bonferroni multiple comparisons test). n = 12/4 (WT) and 10/3 (APP/PS1) recordings/mice. (D) Schematic of the conditioned place preference (CPP) paradigm, including pre-conditioning, conditioning, and post-conditioning phases. (E) Pre-conditioning grid preference showing increased baseline preference for the small grid in APP/PS1 mice compared with WT (two-way ANOVA). (F) CPP test showing increased time spent in the reward-associated context (Cs+) compared with the non-rewarded context (Cs−) in both genotypes, with with increased preference in APP/PS1 mice (two-way ANOVA followed by Bonferroni multiple comparisons test). (G) Chocolate consumption during conditioning showing increased intake in APP/PS1 mice across days (two-way ANOVA). (H) Pellet consumption during conditioning showing no differences between genotypes. Data are presented as mean ± SEM. n = 8 mice per genotype. *p < 0.05, ***p < 0.001.

**Figure 7.**
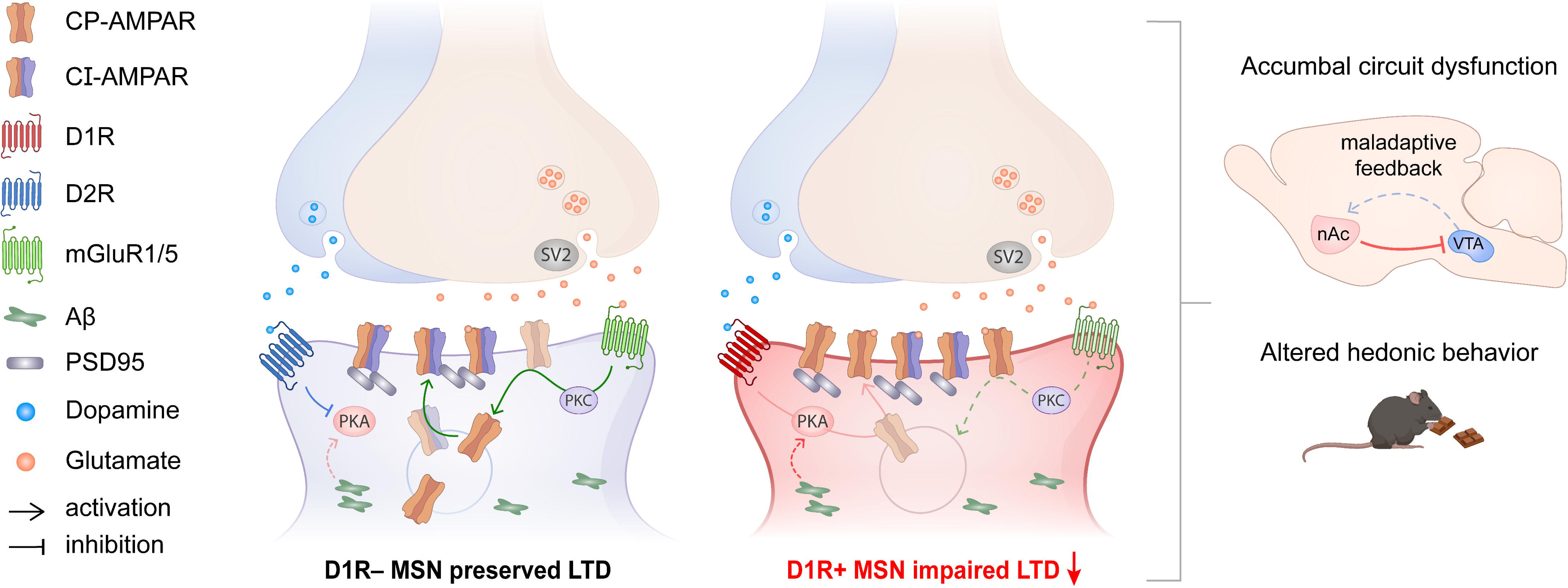
Proposed model of cell-type-specific synaptic alterations in the nucleus accumbens during early stages of Alzheimer’s disease. Schematic representation of glutamatergic and dopaminergic signaling onto accumbal MSNs during pre-plaque stages, characterized by intraneuronal Aβ accumulation in the absence of extracellular plaques. In D1R− MSNs (left; putatively D2R-expressing), synaptic plasticity is preserved, consistent with intact mGluR1/5-dependent LTD. In contrast, D1R+ MSNs (right) exhibit impaired mGluR-dependent LTD, associated with altered AMPAR regulation and increased contribution of NASPM-sensitive AMPAR components. Reduced dopaminergic signaling may contribute to these alterations by modifying D1R-dependent pathways, potentially influencing AMPAR trafficking and synaptic stability. At the circuit level, these cell-type-specific changes are proposed to bias accumbal output and contribute to alterations in reward-related behavior observed during early stages of AD.

We next examined whether this alteration is associated with changes in reward-related behavior. In a chocolate-based conditioned place preference (CPP) paradigm (Fig. 6D), APP/PS1 mice exhibited a baseline preference for the small-grid context, whereas WT mice showed no preference (Fig. 6E). During testing, both genotypes spent more time in the reward-associated context compared with the non-rewarded context (Fig. 6F), indicating intact associative learning. However, APP/PS1 mice consumed more chocolate during conditioning sessions than WT mice (Fig. 6G), while standard pellet consumption did not differ between genotypes (Fig. 6H), suggesting selective alterations in reward-related behavior. No differences were observed between genotypes in anxiety-like behavior or social interaction parameters (Fig. S5A–M), indicating that these behavioral domains remain preserved at this stage.

Together, these results indicate that early alterations in dopaminergic signaling in the nAc of APP/PS1 mice are associated with selective changes in reward-related behavior, while anxiety-like and social behaviors remain preserved. These findings support a domain-specific vulnerability of reward-related processes during pre-plaque stages.

## DISCUSSION

Our study identifies a cell-type-specific disruption of synaptic plasticity in the nAc during early stages of AD pathology. We show that long-term depression is selectively impaired in accumbal D1R+ MSNs despite comparable intracellular Aβ accumulation across neuronal subtypes. This deficit is accompanied by an increased contribution of calcium-permeable AMPARs and reduced dopaminergic signaling, indicating a shift in postsynaptic synaptic gain within accumbal circuits. These alterations occur during pre-plaque stages and are associated with selective changes in reward-related behavior. Together, these findings define an early and cell-type-specific vulnerability of D1R+ MSNs and establish a mechanistic link between intracellular Aβ accumulation and synaptic dysfunction in mesolimbic circuits.

Early dysfunction of the nAc has been reported during early stages of Alzheimer’s disease, including alterations in neuronal activity and network hyperexcitability, yet the synaptic mechanisms underlying these changes remain poorly defined. Here, we identify a postsynaptic mechanism characterized by an increased contribution of calcium-permeable AMPARs together with a selective impairment of mGluR1/5-dependent LTD in D1R+ MSNs. As this form of plasticity normally promotes AMPAR endocytosis and constrains excitatory synaptic strength, its disruption is consistent with the persistence of CP-AMPAR–mediated signaling and a shift toward increased postsynaptic gain. This interpretation is consistent with clinical evidence indicating that neuronal hyperexcitability and epileptiform activity are highly prevalent in patients with AD, emerging early in the disease course and contributing to accelerated cognitive decline (Vossel et al., 2026). Recent computational and neuroimaging approaches have identified a progressive disruption of excitation–inhibition balance across AD stages, with limbic regions showing early vulnerability. Notably, increased excitability within the nAc has been reported at the stage of mild cognitive impairment, preceding overt dementia (Li et al., 2025). These observations are consistent with our findings and support the notion that early synaptic alterations at the cellular level may scale up to circuit-wide dysregulation.

AD is increasingly recognized as a disorder that perturbs distributed neural circuits before the onset of memory impairment, with early non-cognitive symptoms linked to dysfunction in limbic and reward-related networks (Frank et al., 2025; Masters, 2015). In this context, the nAc functions as a central integrative hub where glutamatergic and dopaminergic inputs converge to regulate motivational behavior (Bayassi-Jakowicka et al., 2022). Notably, structural and functional alterations of the nAc have been reported in patients, including reduced volume and disrupted connectivity with prefrontal regions involved in decision-making and inhibitory control (Contreras et al., 2020; Nie et al., 2017). By focusing on pre-plaque stages, our findings provide a mechanistic framework linking early intracellular Aβ accumulation to selective disruption of synaptic plasticity and neuromodulatory control in this circuit. These results extend current models of AD by indicating that early pathology involves mesolimbic dysfunction that cannot be fully explained by hippocampal and cortical alterations alone.

A critical factor in interpreting Alzheimer’s disease mechanisms is the disease stage at which pathology is examined. While most experimental studies have focused on advanced phases characterized by extracellular amyloid plaque deposition and overt cognitive impairment, our findings emphasize an earlier stage in which synaptic and circuit-level alterations are already present in the absence of plaques. In this context, the selective impairment of LTD together with increased CP-AMPAR contribution and reduced dopaminergic signaling supports the notion that early pathology involves a shift in synaptic balance that precedes classical neurodegenerative features. Rather than reflecting late-stage synaptic loss, these changes indicate an initial phase of circuit dysregulation characterized by increased accumbal postsynaptic synaptic gain and reduced local regulatory control. This early synaptic configuration is consistent with growing evidence that intracellular Aβ dynamics play a central role during initial disease stages. Recent work has shown that dendrite-to-dendrite nanotubular networks can mediate the propagation of intracellular Aβ between neurons prior to plaque formation, supporting a model in which early pathology is spatially distributed and not restricted to extracellular aggregation (Chang et al., 2025). Within this framework, our findings extend these observations by demonstrating that such early intracellular pathology is associated with selective disruption of synaptic plasticity and neuromodulatory signaling in the nAc. Over time, persistent disruption of these mechanisms may contribute to maladaptive circuit states and eventual synaptic failure, providing a potential link between early hyperexcitability and later hypoactivity observed in AD progression.

The selective vulnerability of D1R+ MSNs observed in this study parallels findings from addiction and food restriction murine models, in which synaptic plasticity and AMPAR remodeling are preferentially disrupted in D1R+ MSNs within the nAc, linked with mGluR-dependent synaptic plasticity dysfunction (Hwang et al., 2025; Italia et al., 2025; Kawa et al., 2022). Notably, our experiments showed that pharmacological blockade of CP-AMPARs restored synaptic depression, indicating that their retention interferes with LTD expression rather than initiating it. These findings align with forms of synaptic plasticity dysfunction described in other pathological contexts. Together, these results emphasize that the expression of mGluR1/5-dependent plasticity deficits in AD is strongly shaped by circuit context, disease stage, and neuronal subtype.

The selective vulnerability of D1R+ MSNs is likely related to their strong dependence on dopaminergic tone. D1 receptors exhibit lower affinity for dopamine than D2 receptors, rendering D1R+ MSNs particularly sensitive to reductions in dopamine availability (Gerlach et al., 2003; Richfield et al., 1989). Consistent with this framework, we observed reduced dopamine-dependent signaling in the nAc at 6 months of age using a genetically encoded dopamine sensor. These findings align with previous reports showing that dopamine release in the nAc is already diminished during pre-plaque stages in APPswe mice, together with compensatory changes in dopamine transporter expression (Nobili et al., 2017). Although the present data do not allow discrimination between impaired presynaptic release, dopaminergic terminal dysfunction, or early alterations in the integrity of dopaminergic VTA neurons, a reduction in accumbal dopamine availability would be expected to disproportionately weaken D1R-mediated signaling.

Such dopaminergic hypoactivity may converge with Aβ-driven postsynaptic alterations to destabilize the reward direct pathway function during early stages of AD. In support of this idea, Whitcomb et al. demonstrated that intracellular perfusion of Aβ oligomers into hippocampal neurons rapidly increases surface GluA1 expression and promotes CP-AMPAR insertion through a PKA-dependent mechanism (Whitcomb et al., 2015). A reduction in dopaminergic tone may therefore weaken physiological D1R–PKA coupling, generating conditions in which intraneuronal Aβ aberrantly engages PKA signaling to drive GluA1 membrane insertion. In contrast, D2R signaling suppresses adenylyl cyclase activity and limits PKA activation (Beaulieu and Gainetdinov, 2011), potentially restricting this mechanism in D1R− MSNs and contributing to their relative resilience. Together, impaired mGluR1/5-dependent AMPAR endocytosis and Aβ-associated GluA1 insertion would bias synapses toward persistent CP-AMPAR enrichment, providing a mechanistic framework for the selective loss of LTD observed in D1R+ MSNs during early AD.

In agreement with the present findings, Aguado et al. reported increased CP-AMPAR expression in the hippocampus of cognitive vulnerable APPswe mice at advanced disease stages (Aguado et al., 2024). Notably, mGluR5 expression was selectively reduced in vulnerable APPswe mice but preserved in resilient animals, paralleling the normalization of CP-AMPAR levels (Aguado et al., 2024). These observations suggest that coordinated regulation of mGluR1/5 signaling and CP-AMPAR composition may modulate synaptic vulnerability across disease stages and brain regions. Complementing this view, Guo et al. demonstrated that acute exposure to exogenous oligomeric Aβ in the nAc of young WT mice induces synaptic incorporation of CP-AMPARs, leading to spine loss, synaptic weakening, and motivational deficits (Guo et al., 2022). Importantly, this model reflects extracellular Aβ-driven pathology and preferentially affects D2 MSNs, in contrast to our findings showing that endogenous intraneuronal Aβ accumulation during early stages is associated with selective CP-AMPAR incorporation in D1R+ MSNs. Together, these studies support CP-AMPAR dysregulation as a convergent mechanism contributing to synaptic dysfunction across disease stages, while emphasizing that the cellular context and subcellular localization of Aβ critically shape its synaptic consequences.

The nAc functions as a key inhibitory hub within the mesolimbic circuit, regulating reward-related signal gain through the integration of glutamatergic inputs and dopaminergic modulation (Russo and Nestler, 2013). The selective loss of LTD in D1R+ MSNs is therefore expected to bias circuit output toward enhanced direct pathway activity, reducing the ability of the nAc to constrain excitatory drive and favoring heightened reward-seeking responses. In parallel, reduced dopaminergic tone may further attenuate D1R signaling, potentially promoting compensatory increases in reward consumption to achieve comparable motivational salience. Consistent with this framework, both genotypes exhibited comparable conditioned place preference, indicating preserved associative learning and contextual preference, and no differences were detected in anxiety-like behavior or social interaction. However, APP/PS1 mice displayed a selective increase in consumption of palatable solid food, while intake of standard chow remained unchanged. Together, these findings indicate that early synaptic dysfunction within the nAc is associated with altered hedonic valuation rather than deficits in reward learning or broader affective behaviors.

Some limitations should be considered when interpreting these findings. First, all experiments were conducted exclusively in male mice, precluding assessment of sex-specific mechanisms. This is relevant given evidence that estradiol signaling modulates synaptic plasticity in the nAc through mGluR5- and endocannabinoid-dependent pathways and influences reward circuit function (Peterson et al., 2016). Moreover, synaptic plasticity in this region can rely on distinct molecular mechanisms in males and females despite similar functional outcomes, as shown by differences in the signaling pathways underlying long-term potentiation (Copenhaver and LeGates, 2024). Second, although our data reveal a strong association between intraneuronal Aβ accumulation and synaptic alterations, the APP/PS1 model does not allow definitive attribution of these effects exclusively to Aβ. Nonetheless, converging evidence supports a direct role for intraneuronal Aβ in modulating excitatory synaptic function. Intraneuronal infusion of human and synthetic Aβ oligomers has been shown to enhance neuronal synchronization and AMPAR-mediated transmission (Fernandez-Perez et al., 2021), and to increase CP-AMPAR–mediated currents in cultured nAc MSNs derived from APP/PS1 mice (Saavedra-Sieyes et al., 2025). Together, these findings support a role for intraneuronal Aβ in shaping excitatory synaptic function, specifically through its association with altered AMPAR composition and impaired LTD in D1R+ MSNs. While the present data do not establish direct causality, they indicate that intraneuronal Aβ accumulation is closely linked to the synaptic mechanisms underlying CP-AMPAR enrichment and plasticity deficits observed in the nAc. Future studies incorporating sex as a biological variable and experimental strategies enabling selective manipulation of intraneuronal Aβ will be required to determine causal relationships.

In conclusion, this study identifies early synaptic alterations in the nAc characterized by disrupted AMPAR composition, selective impairment of LTD in D1R+ MSNs, and reduced dopaminergic signaling prior to plaque formation. These changes are accompanied by selective alterations in reward-related behavior, supporting the view that early non-cognitive symptoms arise from circuit-level imbalance rather than late-stage neurodegeneration. Together, these findings position the nAc as an early site of vulnerability and identify neuronal subtype identity as a key determinant of synaptic dysfunction in AD, providing a mechanistic framework linking intraneuronal Aβ accumulation to early mesolimbic circuit dysregulation and highlighting CP-AMPAR signaling and mGluR1/5-dependent plasticity as candidate pathways for therapeutic modulation at early disease stages.

## Acknowledgements

We thank Laurie Aguayo, Helena Zambrano, and Ailine Riquelme for technical assistance, as well as Jocelyn González and Ixia Cid for support during experimental procedures. We also acknowledge Carolina Benítez (CREAV-UDEC) and Claudia Ramírez for veterinary assistance. We thank Lauren Aguayo and Mauricio Avendaño Valenzuela (Universidad de Concepción, Chile) for assistance with language editing.

## Author contributions

N.R.L.: Conceptualization, methodology, investigation, formal analysis, data curation, visualization, writing – original draft, writing – review & editing. Designed all main figures (Figs. 1–7) and the graphical abstract. Performed all experiments and data analyses for Figs. 1–4, 7 and Supplementary Figs. S1, S2, and S4. Supervised and validated data corresponding to experiments not directly performed by the author. I.M.: Investigation (Supplementary Fig. S4 recordings), genotyping, data acquisition and analysis (Fig. 1), writing – review & editing. J.G.S.: Investigation (Supplementary Fig. S5), data interpretation, writing – review & editing. P.S.S.: Investigation and formal analysis (Supplementary Fig. S3), writing – review & editing. L.A.W.: Investigation and formal analysis (Fig. 6), writing – review & editing, funding acquisition. A.S.: Methodology, supervision of GCaMP and dLight photometric recordings, data interpretation, writing – review & editing. L.S.M.: Conceptualization, supervision, writing – review & editing, funding acquisition. L.G.A.: Conceptualization, supervision, writing – review & editing, project administration, funding acquisition.

## Data availability

The data supporting the findings of this study are available from the corresponding author upon reasonable request.

---

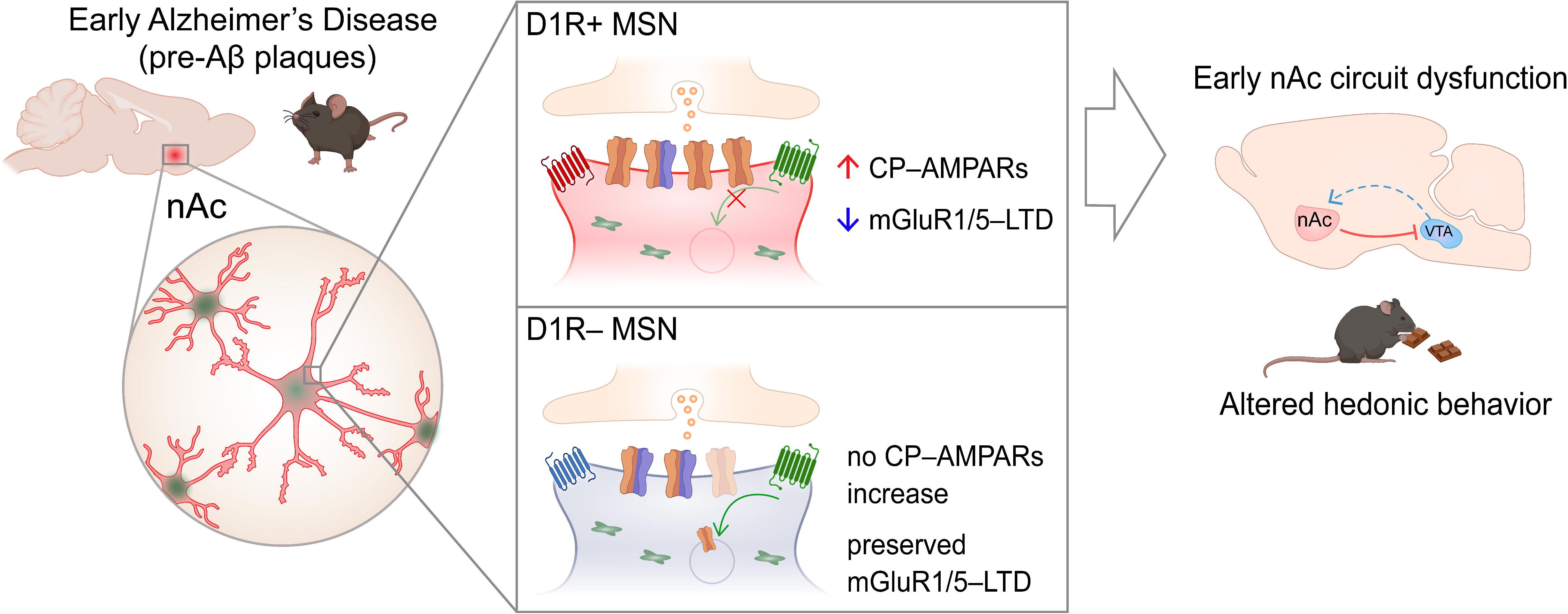

