## Supplemental figures 1 - 5 for "Selective impairment of long-term depression in accumbal D1R+ MSNs involves calcium-permeable AMPARs in early Alzheimer’s disease"

**Supplementary Figure 1**

**Supplementary Figure 2**

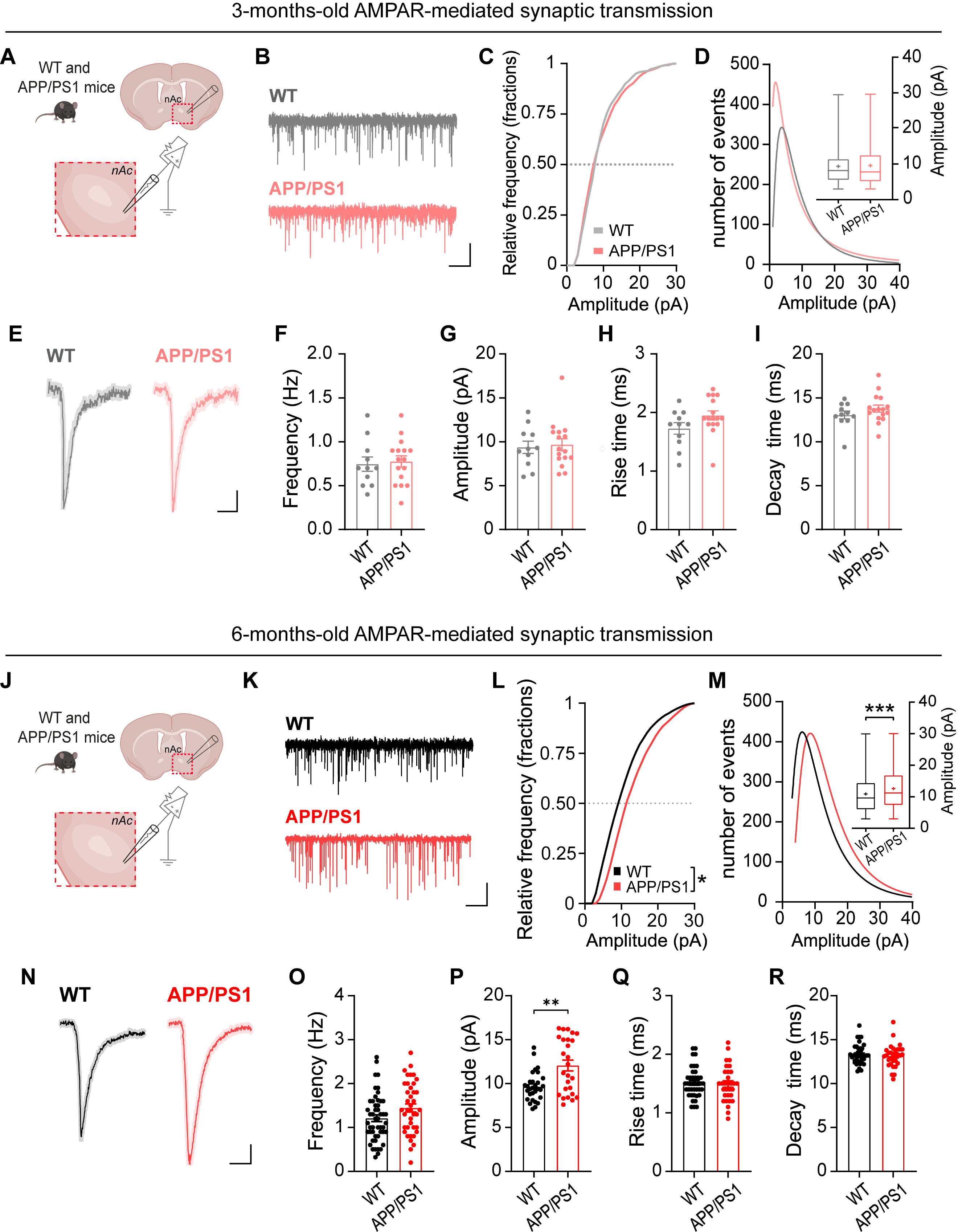

**Supplementary Figure 3**

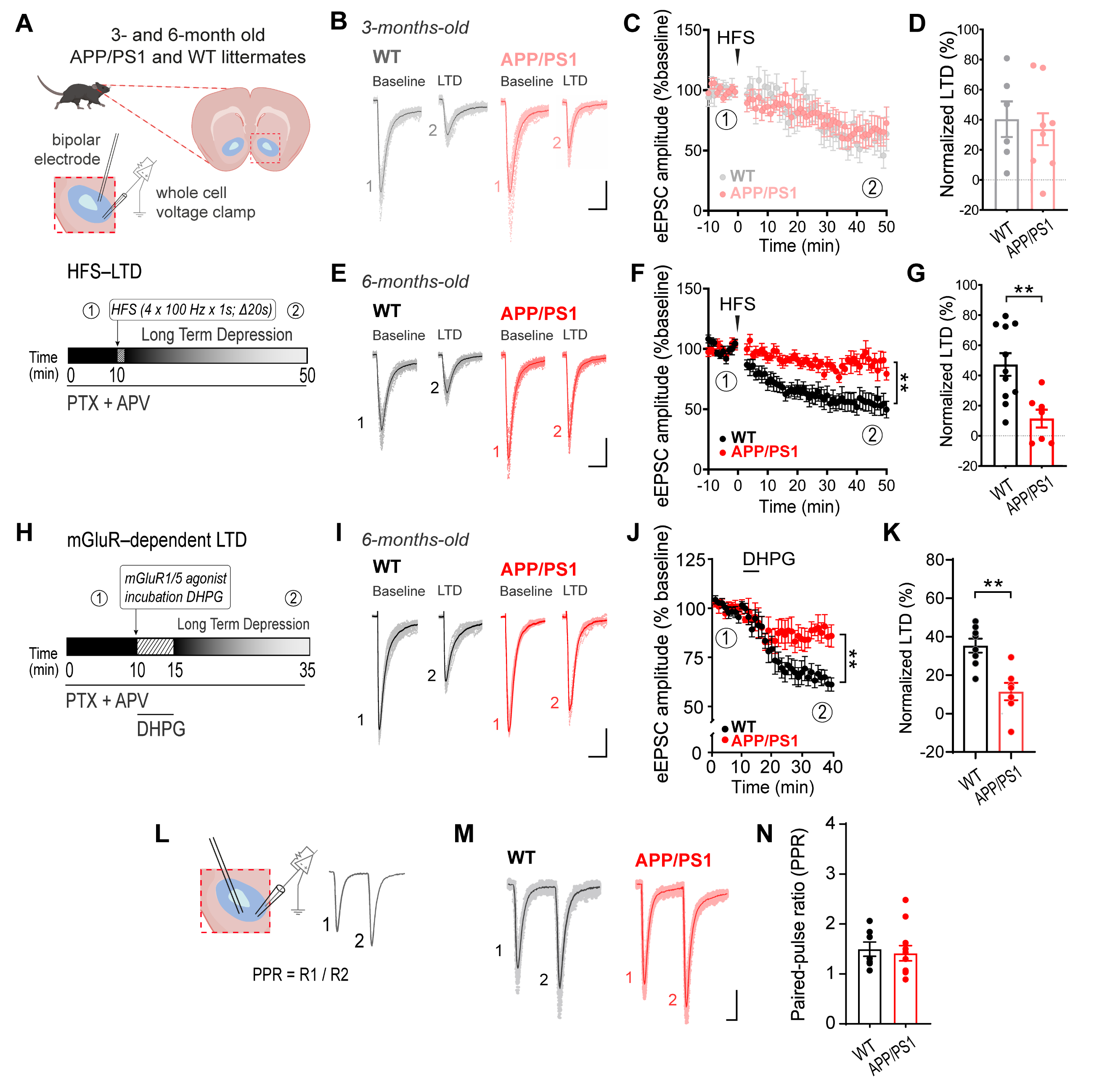

**Supplementary Figure 4**

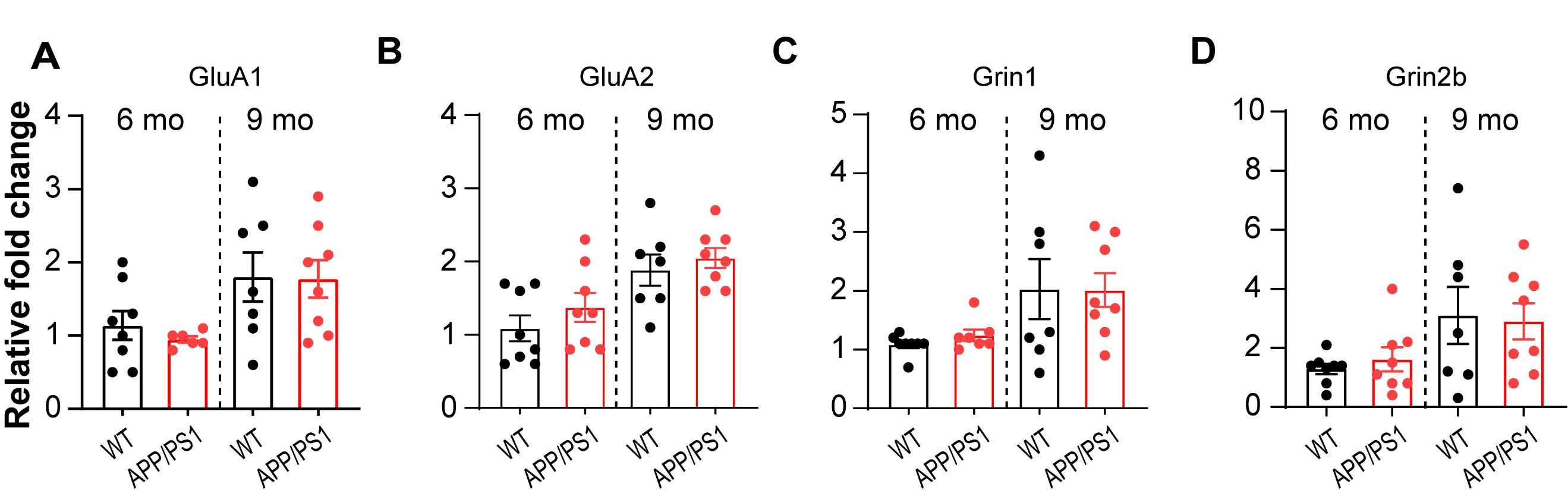

**Supplementary Figure 5**

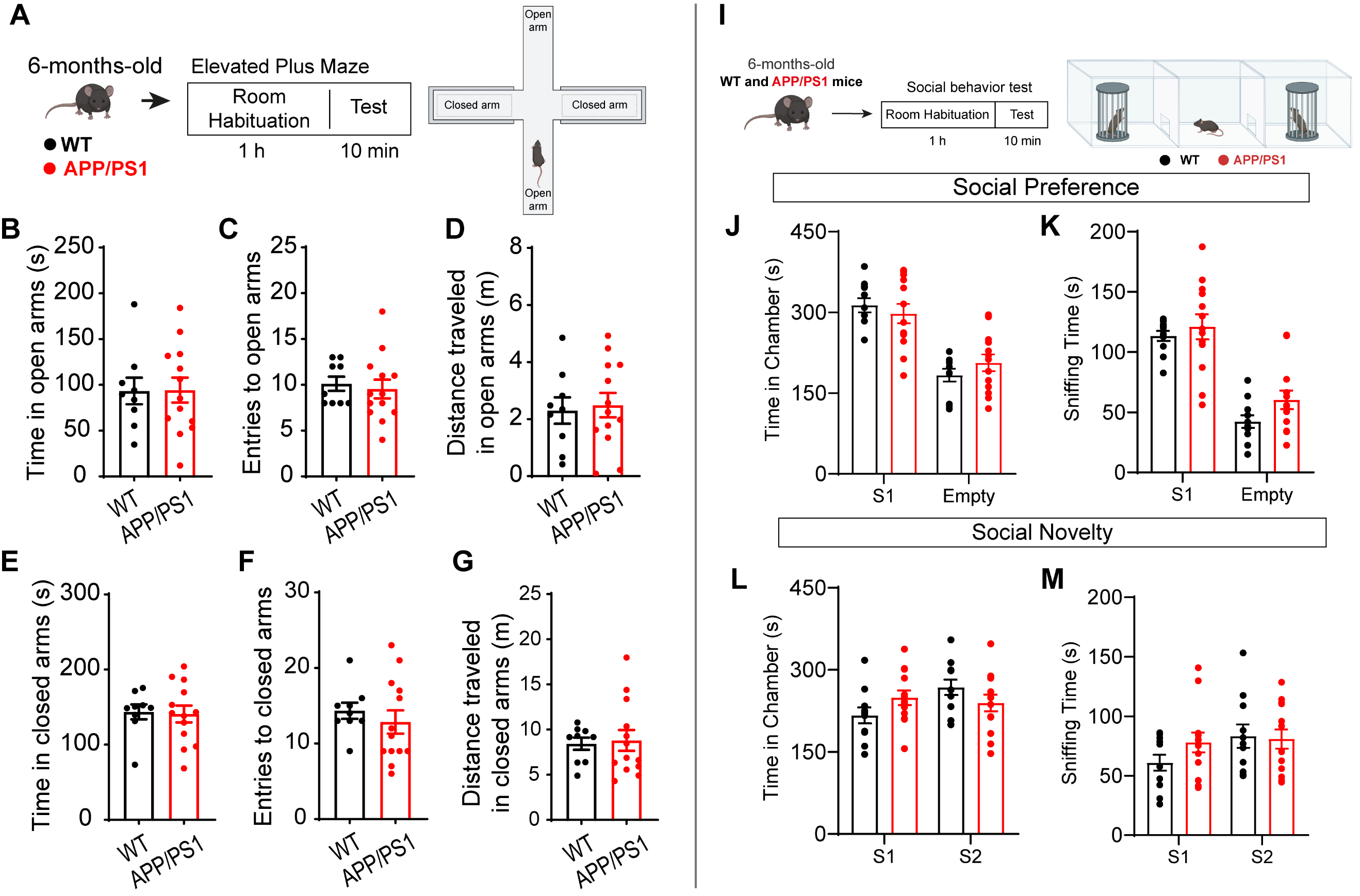

**Supplementary Figures legends**

**Fig. 1 Supplementary.** Intracellular Aβ accumulation, absence of extracellular plaque deposition, and preserved microglial profile in the nAc during pre-plaque stages in APP/PS1 mice. (A) Representative immunohistochemistry of coronal nAc sections (30 μm) from 3-month-old WT and APP/PS1 mice labeled with MOAβ-2 (green), MAP2 (magenta), and Hoechst (blue). APP/PS1 mice show increased intracellular Aβ immunoreactivity in neuronal somata. Insets (dashed boxes) illustrate higher magnification views of individual neurons. Scale bars: 50 µm. (B) Quantification of MOAβ-2 fluorescence intensity measured within neuronal somata (MAP2 mask) reveals significantly higher intracellular Aβ levels in APP/PS1 mice compared with WT (unpaired two-tailed t-test, ***p < 0.001). Each point represents the mean value per animal (WT n = 5; APP/PS1 n = 5), with 20 neurons analyzed per animal. (C) Representative Thioflavin-S (Th-S) staining in coronal sections from WT and APP/PS1 mice at 6 and 12 months of age in the nAc (top) and cortex (bottom). No detectable Th-S–positive signal is observed in the nAc at 6 months in either genotype, whereas sparse plaque deposition is evident at 12 months in APP/PS1 mice (arrowheads). In contrast, cortical plaques are detectable at earlier stages and increase with age. Scale bars: 50 µm. (D) Representative immunohistochemistry of nAc sections from 6-month-old WT and APP/PS1 mice labeled with Iba1 (green), MAP2 (magenta), tdTomato (red), and Hoechst (blue). Merged images show comparable microglial distribution across genotypes. Scale bars: 50 µm. (E) Quantification of Iba1 fluorescence intensity in the nAc shows no significant differences between WT and APP/PS1 mice (unpaired two-tailed t-test, ns). Each point represents a single z-projection image (30 μm stack; field size 581 × 581 µm) obtained from 2–3 slices per animal (WT n = 4; APP/PS1 n = 4). (F) Quantification of Iba1+ cell density (cells/mm²) in the nAc shows no differences between genotypes (Welch’s t-test, ns). Each point represents the mean per animal calculated from multiple ROIs (4 ROIs per image; ≥6 images from ≥3 slices per animal; WT n = 4; APP/PS1 n = 4).

**Fig. 2 Supplementary.** Age-dependent alterations in AMPAR-mediated excitatory synaptic transmission in the nucleus accumbens of APP/PS1 mice. (A–I) Analysis of AMPAR-mediated spontaneous excitatory postsynaptic currents (sEPSCs) recorded from medium spiny neurons (MSNs) in the nucleus accumbens (nAc) of 3-month-old WT and APP/PS1 mice. (A) Schematic representation of the experimental configuration for whole-cell voltage-clamp recordings of sEPSCs in acute nAc slices. (B) Representative sEPSC traces recorded at −60 mV from WT (grey) and APP/PS1 (light red) MSNs. (C) Cumulative probability distribution of sEPSC amplitudes showing overlapping distributions between WT and APP/PS1 MSNs (P = 0.837, Mann–Whitney test). (D) Amplitude distribution histogram with box plot summarizing sEPSC amplitudes, showing no significant difference between genotypes (median WT = 8.16 pA, APP/PS1 = 7.82 pA; P = 0.837, Mann–Whitney test; geometric mean WT = 7.37 pA, APP/PS1 = 6.47 pA). (E) Representative averaged sEPSC events from WT and APP/PS1 MSNs. F Quantification of sEPSC frequency showing no significant differences between genotypes (median WT = 0.70 Hz, APP/PS1 = 0.80 Hz; P = 0.619, Mann–Whitney test). (G) Quantification of sEPSC amplitude showing no significant differences (median WT = 9.20 pA, APP/PS1 = 9.05 pA; P = 0.952, Mann–Whitney test). (H) Rise time analysis showing no significant differences between genotypes (median WT = 1.80 ms, APP/PS1 = 1.90 ms; P = 0.092, Mann–Whitney test). (I) Decay time analysis showing no significant differences (mean WT = 13.05 ms, APP/PS1 = 13.78 ms; P = 0.250, unpaired two-tailed t-test). (J–R) Analysis of AMPAR-mediated sEPSCs recorded from MSNs in the nAc of 6-month-old WT and APP/PS1 mice. (J Schematic representation of the experimental configuration for whole-cell voltage-clamp recordings in 6-month-old mice. (K) Representative sEPSC traces recorded at −60 mV from WT (black) and APP/PS1 (red) MSNs. (L) Cumulative probability distribution of sEPSC amplitudes showing a rightward shift in APP/PS1 MSNs compared with WT (P < 0.05, Mann–Whitney test). (M) Amplitude distribution histogram with box plot showing a significant increase in mean sEPSC amplitude in APP/PS1 MSNs compared with WT (Welch’s t-test, P < 0.001; geometric mean WT = 9.32 pA, APP/PS1 = 10.95 pA; ratio = 1.18, 95% CI = 1.15–1.20; P = 0.0208). (N) Representative averaged sEPSC events from WT and APP/PS1 MSNs. (O) Quantification of sEPSC frequency showing no significant differences between genotypes (median WT = 1.15 Hz, APP/PS1 = 1.40 Hz; P = 0.052, Mann–Whitney test). (P) Quantification of sEPSC amplitude showing a significant increase in APP/PS1 MSNs compared with WT (median WT = 9.45 pA, APP/PS1 = 11.70 pA; P = 0.006, Mann–Whitney test). (Q) Rise time analysis showing no significant differences between genotypes. (R) Decay time analysis showing no significant differences between genotypes. Each data point represents a single recorded neuron. Data are presented as mean ± SEM unless otherwise indicated. Statistical tests and exact P values are reported for each panel.

**Fig. 3 Supplementary.** Age-dependent impairment of synaptic depression mechanisms in the nucleus accumbens of APP/PS1 mice. (A) Schematic representation of the experimental design for whole-cell voltage-clamp recordings of electrically evoked EPSCs from MSNs in the nAc of 3- and 6-month-old WT and APP/PS1 mice. The experimental timeline indicates baseline acquisition (①) and quantification of LTD expression (②), calculated as the mean eEPSC amplitude during the final 5 min of recording normalized to the 10-min baseline period. (B) Representative average traces of AMPAR-mediated eEPSCs recorded from MSNs of 3-month-old WT (grey) and APP/PS1 (light red) mice during baseline and after HFS-induced LTD. Overlaid traces are shown for comparison. (C) Time course of normalized eEPSC amplitude in 3-month-old mice showing comparable HFS-LTD expression in WT and APP/PS1 MSNs. (D) Quantification of normalized LTD magnitude (%) at 3 months reveals no significant difference between genotypes (unpaired two-tailed t-test, t(12) = 0.4158, P = 0.6849). (E) Representative average eEPSC traces recorded from MSNs of 6-month-old WT (black) and APP/PS1 (red) mice during baseline and after HFS-LTD induction. (F) Time course of normalized eEPSC amplitude in 6-month-old mice shows reduced HFS-LTD expression in APP/PS1 MSNs compared with WT. (G) Quantification of normalized LTD magnitude (%) at 6 months demonstrates a significant reduction of HFS-LTD in APP/PS1 MSNs relative to WT (unpaired two-tailed t-test, t(16) = 3.386, P = 0.0038). (H) Schematic representation of whole-cell voltage-clamp recordings in nAc MSNs from 6-month-old WT and APP/PS1 mice during chemical LTD induction. (I) Representative average eEPSC traces recorded before and after DHPG application (50 μM) in WT and APP/PS1 MSNs. (J Time course of normalized eEPSC amplitude following DHPG application shows robust mGluR1/5-dependent LTD in WT MSNs but not in APP/PS1 MSNs. (K) Quantification of DHPG-induced LTD reveals a significant impairment in APP/PS1 MSNs compared with WT. Statistical analysis was performed using an unpaired two-tailed t-test, showing a significant reduction in LTD magnitude in APP/PS1 neurons (t(13) = 4.155, P = 0.0011; WT mean = 35.33%, APP/PS1 mean = 11.45%). (L) Schematic illustration of the paired-pulse stimulation protocol used to assess presynaptic release probability, with paired-eEPSCs evoked at a fixed interstimulus interval and the PPR, calculated as the ratio of the second to the first response (R/R’). (M) Representative average paired-eEPSC recordings from MSNs in the nAc of WT (black) and APP/PS1 (red) mice. (N) Quantification of PPR shows no significant difference between genotypes (unpaired two-tailed t-test, t(16) = 0.364, P = 0.721), indicating preserved presynaptic release probability. Data are shown as mean ± s.e.m., and each data point represents a single recorded neuron. Sample sizes: 3-month-old HFS-LTD, WT n = 6 neurons (3 mice), APP/PS1 n = 8 neurons (3 mice); 6-month-old HFS-LTD, WT n = 11 neurons (5 mice), APP/PS1 n = 7 neurons (4 mice); DHPG-LTD, WT n = 8 neurons (4 mice), APP/PS1 n = 7 neurons (4 mice); PPR analysis, WT n = 11 neurons (3 mice), APP/PS1 n = 7 neurons (3 mice). **P < 0.01

**Fig. 4 Supplementary.** Relative expression of glutamate receptor subunits in the nucleus accumbens of APP/PS1 mice. (A – D) Relative mRNA expression levels assessed by RT–qPCR for Gria1 (GluA1), Gria2 (GluA2), Grin1 (GluN1), and Grin2b (GluN2B) in the nucleus accumbens of WT and APP/PS1 mice at the indicated ages. No significant differences were detected between genotypes for any transcript at any time point analyzed (unpaired two-tailed t-tests or two-way ANOVA where appropriate; all P > 0.05). Data are shown as mean ± s.e.m. Each data point represents one animal.

**Fig. 5 Supplementary.** Anxiety-like behavior and social interaction are preserved in 6-month-old APP/PS1 mice. (A) Experimental design of the elevated plus maze test performed in 6-month-old WT and APP/PS1 mice, including a room habituation phase followed by a 10 min test session. (B–D) Quantification of time spent in the open arms, number of entries into the open arms, and distance traveled in the open arms during the elevated plus maze test. (E–G) Quantification of time spent in the closed arms, number of entries into the closed arms, and distance traveled in the closed arms. No genotype-dependent differences were observed across any elevated plus maze parameter. Bars represent mean ± SEM; WT n = 11 mice and APP/PS1 n = 13 mice. I Experimental design of the social interaction test performed in 6-month-old WT and APP/PS1 mice, including room habituation followed by a 10 min test session. (J–K) Social preference phase showing time spent in each chamber and sniffing time directed toward a conspecific mouse compared with an empty cage. (L–M) Social novelty phase showing time spent in each chamber and sniffing time directed toward a familiar mouse versus a novel mouse. No significant genotype-dependent differences were detected in social preference or social novelty measures. Bars represent mean ± SEM; WT n = 9 mice and APP/PS1 n = 13 mice.
